# rvTWAS: identifying gene-trait association using sequences by utilizing transcriptome-directed feature selection

**DOI:** 10.1101/2023.07.16.549227

**Authors:** Jingni He, Qing Li, Qingrun Zhang

## Abstract

Towards the identification of genetic basis of complex traits, transcriptome-wide association study (TWAS) is successful in integrating transcriptome data. However, TWAS is only applicable for common variants, excluding rare variants in exome or whole genome sequences. This is partly because of the inherent limitation of TWAS protocols that rely on predicting gene expressions. Briefly, a typical TWAS protocol has two steps: it trains an expression prediction model in a reference dataset containing gene expressions and genotype, and then applies this prediction model to a genotype-phenotype dataset to “impute” the unobserved expression (that is called GReX) to be associated to the phenotype. In this procedure, rare variants are not used due to its low power in predicting expressions. Our previous research has revealed the insight into TWAS: the two steps are essentially genetic feature selection and aggregations that do not have to involve predictions. Based on this insight disentangling TWAS, rare variants’ inability of predicting expression traits is no longer an obstacle. Herein, we developed “rare variant TWAS”, or rvTWAS, that first uses a Bayesian model to conduct expression-directed feature selection and then use a kernel machine to carry out feature aggregation, forming a model leveraging expressions for association mapping including rare variants. We demonstrated the performance of rvTWAS by thorough simulations and real data analysis in three psychiatric disorders, namely schizophrenia, bipolar disorder, and autism spectrum disorder. rvTWAS will open a door for sequence-based association mappings integrating gene expressions.

## INTRODUCTION

Integrating transcriptome in genotype-phenotype association studies, transcriptome-wide association study (TWAS) represents a successful example in utilizing-omics data in gene mapping (Gamazon *et al*. 2015; Gusev *et al*. 2016). A typical TWAS tool is implemented as a two-step protocol: First, for a gene, one can form an expression prediction model using (usually cis) genotype: e ∼ Σ β_i_X_i[ref]_ + *ε* in a reference dataset [e.g., GTEx (GTEx Consortium 2013; GTEx Consortium 2020)]. The predicted expression, *ê*, is called Genetically Regulated eXpression, or GReX, for this gene. Next, in the second step, in the main dataset for genome-wide association study, or GWAS, (which doesn’t contain expressions), one can predict expression using the genotype: *ê* = Σ β_i_X_i[GWAS]_, and then use the predicted expression *ê* to conduct association mapping *y* ∼ *ê*. The outcome will be whether this gene’s genetic variants are (aggregately) associated with the phenotype *y*. This procedure can be carried out for all genes with good heritability (e.g., >1%), therefore is termed as “transcriptome-wide”. TWAS tools have been widely used in integrating transcriptomes with legacy GWAS data and has led to the discovery of novel genes in many diseases (Gusev *et al*. 2016; Gusev *et al*. 2018; Lu *et al*. 2018; Wu *et al*. 2018; Wu *et al*. 2019; Cao *et al*. 2021b; Guo *et al*. 2021; He *et al*. 2022).

Despite its success in using legacy GWAS data, TWAS protocols do not utilize rare genetic variants. The reason may be that rare variants have little power in predicting gene expressions for their low frequency in the population. In contrast, association mapping using rare genetic variants is an established research field (Lee *et al*. 2014; Auer and Lettre 2015; He *et al*. 2017; Momozawa AND Mizukami 2021), especially since the availability of the high-throughput instruments popularizing whole-exome or whole-genome sequencing. Rare variants account for much of the human mutational catalog (Momozawa AND Mizukami 2021), and their importance in gene mapping is justified by the rationale that disease-causal variants could be under negative selection therefore being rare in the population (Charlesworth *et al*. 1995; O’CONNOR *et al*. 2019; Schoech *et al*. 2019). However, due to the sparsity of the rare variants (i.e., not sufficiently shared across individuals), there is little promise in using them to conduct predictions. As such, no efforts using rare variants for TWAS because expressions are not “predictable” using rare variants.

Our recent works showed that the interpretation of “predicting expressions” did not reveal the essence of TWAS, if not misleading (Cao *et al*. 2021a; Cao *et al*. 2021b; Cao *et al*. 2022). First, theoretical power analysis showed that TWAS could be more powerful than the hypothetical scenario in which expression data is available in the main GWAS dataset (Cao *et al*. 2021a).

Additionally, TWAS could be underpowered comparing with GWAS when the expression heritability is low (Cao *et al*. 2021a). Both results question the interpretation of the prediction of expression in TWAS. As such, we proposed to interpret the “prediction” step as a selection of genetic variants directed by expression. From the perspective of Machine Learning, the first step in TWAS is exactly feature selection and the second being feature aggregation (Cao *et al*. 2022). With this interpretation in mind, one may cancel GReX and instead conduct feature selection and aggregation independently using whatever methods fit best. Indeed, novel methods splitting these two steps developed by us (Cao *et al*. 2021b; Cao *et al*. 2022) and others (Tang *et al*. 2021) showed higher power than standard TWAS.

Our above insight opens a door to developing tools covering broader ranges of data, for instance rare variants in this work. By considering a TWAS protocol as the combination of feature selection (instead of training a prediction) and aggregation (instead of applying a prediction), the instability of predicting expressions using rare variants can be skipped.

Particularly, as long as one can select rare variants associating with gene expressions and aggregate them for downstream genotype-phenotype mapping, transcriptomes can be utilized regardless of whether one could predict them. Indeed, many tools facilitate the above two procedures. As far as our knowledge, Bayesian model selection methods using posterior inclusion probability, or PIP represents state-of-the-art in variant selections (Wang *et al*. 2020). Also, kernel-machine methods (Wu *et al*. 2011) have been widely used in aggregating rare variants in association mapping.

Based upon the above rationale and preliminary development, we propose rare-variant TWAS, or rvTWAS, which is composed by two steps: First, rvTWAS uses Sum of Single Effects, or SuSiE (Wang *et al*. 2020) to carry out variants selections to form a prioritized set of genetic variants (including rare variants) weighted by their relevance to gene expressions (**Figure 1A**). Second, supported by our previous successful attempt using kernel methods to carry out common variants TWAS (Cao *et al*. 2021b; Cao *et al*. 2022), as well as the community’s practice of using kernel models in both common and rare variants GWAS (Wu *et al*. 2011; Lee *et al*. 2012), rvTWAS uses a kernel method (Wu *et al*. 2011; Lee *et al*. 2012) to aggregate weighted variants to form a score test for the association (**Figure 1B**). We carried out thorough simulations using various genetic architecture and causality models to test the performance of rvTWAS. We also applied rvTWAS to real sequencing data for association mapping in schizophrenia (SCZ) (Kiezun *et al*. 2012; Pasaniuc *et al*. 2012), bipolar disorder (BPD) (Kiezun *et al*. 2012; Pasaniuc *et al*. 2012) and autism spectrum disorder (ASD) (Neale *et al*. 2012; DE RUBEIS *et al*. 2014), leading to the discovery of additional genes underlying these psychiatric disorders which have greater enrichment in a comprehensive disease gene database (Pinero *et al*. 2015; Pinero *et al*. 2017; Pinero *et al*. 2020).

**Figure 1.**
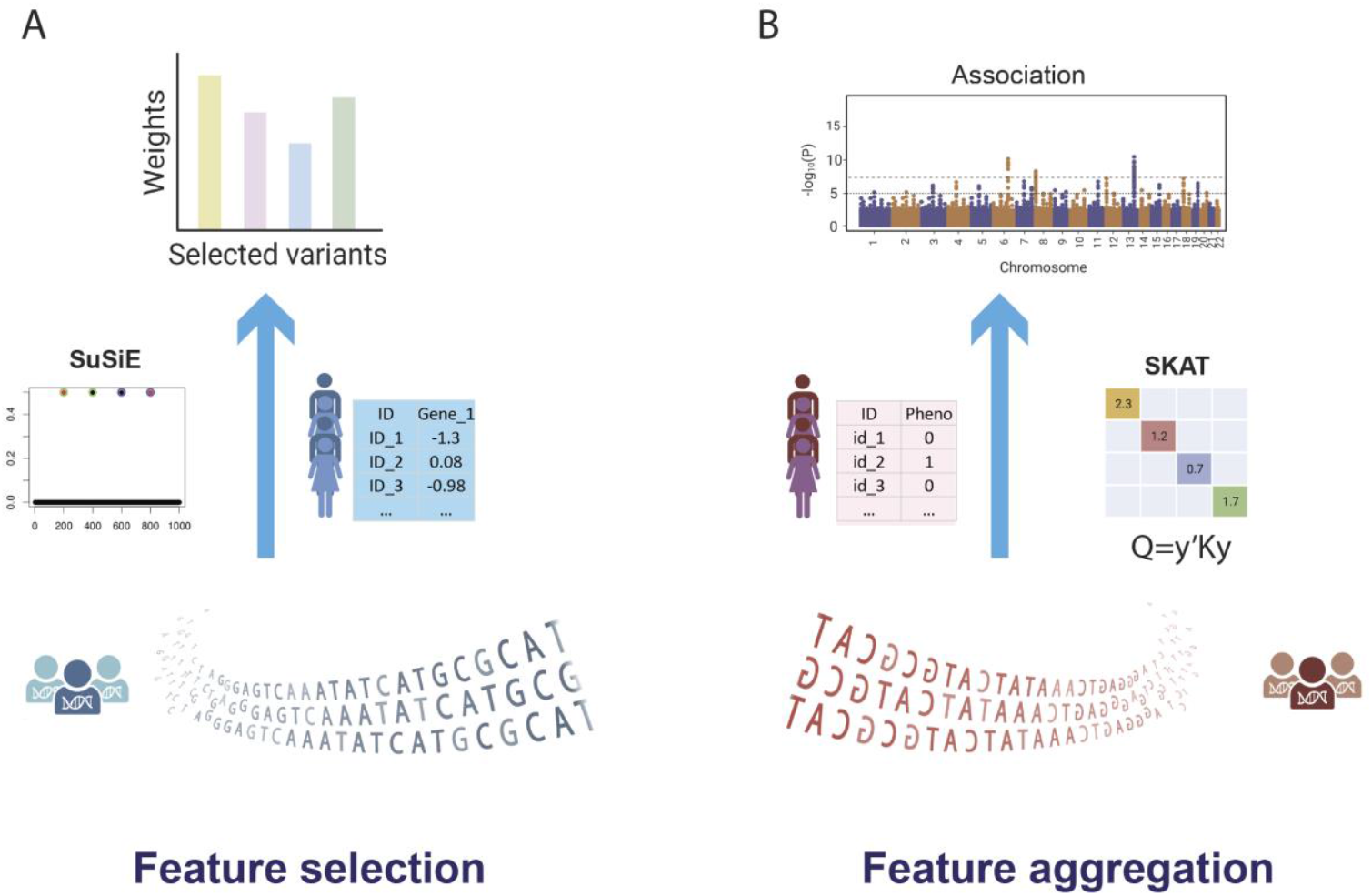
Overview of the rvTWAS framework. **A)** The framework uses a reference gene expression dataset for feature selection by SuSiE. **B)** Weights derived from the feature selection step are then used in feature aggregation in the GWAS dataset. These aggregated features are then associated with the phenotype of interest to identify variants of interest by SKAT.

## MATERIALS AND METHODS

### The rvTWAS Model

As depicted in (**Figure 1**), rvTWAS uses the Bayesian feature selection model implemented by SuSiE (Wang *et al*. 2020) to select variants that are highly associated with gene expressions and aggregates them for association mapping to the phenotype using a weighted kernel (Wu *et al*. 2011; Lee *et al*. 2012). rvTWAS works on one gene each time (and carries out the analysis of all genes sequentially). For each gene, the input data are expressions of this gene and cis genetic variants surrounding the gene (by default, 500Kb of flanking region). One may specify the preferred minor allele frequency (MAF) as the cut-off of qualified variants into the analysis. For instance, one can set up a cut-off = 0.005 (only rare variants with MAF<0.5% are included); however one may specify no cut-off (= all variants included). In Step 1, rvTWAS runs SuSiE using its default parameters, which leads to the outcome of sets of -cis genetic variants, which are termed as “credible sets” by SuSiE. For the credible sets of variants, SuSiE outputs a regression coefficient 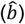 for each variant, indicating its importance quantified by the selected model (**Figure 1A**). By default, rvTWAS includes all variants within all the credible sets. In Step 2, for each gene, by using the regression coefficients 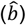 as weights of selected variants, rvTWAS aggregates these variants using a weighted kernel as implemented in Sequence Kernel Association Test, or SKAT (Wu *et al*. 2011; Lee *et al*. 2012) (**Figure 1B**). Briefly speaking, rvTWAS calculates a weighted genomic relationship matrix (GRM) by *K*_*w*_ = *G*^T^*WG*/n (where *G* is the local genotype matrix of selected genetic variants, *W* is the diagonal matrix formed by the weights (i.e., regression coefficient 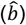 of corresponding variants, and n is the number of variants). Using this GRM, rvTWAS further forms a *Q*-score by calculating *Q* = *y*^′^*K*_*w*_*y*, where *y* is the phenotype (after adjusting co-factors). This *Q*-score follows a mixture chi-squared distribution (Wu *et al*. 2011; Lee *et al*. 2012). By testing the significance of this *Q*-score to all genes, rvTWAS completes the genome-wide association analysis.

As mentioned above, by adjusting the MAF cut-off, rvTWAS supports both rare variants analysis and the inclusive joint analysis of rare and common variants. Our practice showed that the protocol of inclusively analyzing rare and common variants leads to better results. Therefore, the main results (to be presented in **Results**) are based on the inclusive analysis of both rare and common variants, leaving the disentanglement of contributions from common and rare variants in **Discussion**.

### Alternative Methods compared to rvTWAS

In this work, we compared the performance of rvTWAS to multiple alternative methods, namely PrediXcan(Gamazon *et al*. 2015), kTWAS (Cao *et al*. 2021b), and mkTWAS (Cao *et al*. 2022). Here we briefly describe the mathematical models of alternative methods for an intuitive comparison, leaving details to their corresponding publications.

***PrediXcan***(Gamazon *et al*. 2015) is the representative TWAS protocol. We chose this tool to compare rvTWAS to standard TWAS. As described in **Introduction**, PrediXcan first trains an expression prediction model e ∼ Σ β_i_X_i[ref]_ + *ε* in a reference dataset (e.g., GTEx (GTEx Consortium 2013; GTEx Consortium 2020)) using cis genetic variants (where e is the gene expression, X_i[ref]_ is the *i*-th genetic variant in the region, and β_i_ is the coefficient to be estimated). Next, in the GWAS dataset, using the coordinate-matched genotype X_i[GWAS]_, it predicts expression: *ê* = Σ β_i_X_i[GWAS]_, and then use the predicted expression *ê* to conduct association mapping *y* ∼ *ê* (where *y* is the phenotype). Particularly, the variants within the +/-500Kb flanking region of the gene are included in the analysis, and Elastic Net (Zou AND Hastie 2005) is used to train the coefficients β_i_.

***kTWAS*** (kernel-TWAS) (Cao *et al*. 2021b), and ***mkTWAS*** (marginal kernel-TWAS) (Cao *et al*. 2022) are our tools using a kernel machine to replace GReX for feature aggregation in a TWAS protocol. We have shown these tools outperform standard TWAS when analyzing common variants (Cao *et al*. 2021b; Cao *et al*. 2022), which is in principle echoed by others using a similar tool(Tang *et al*. 2021). We choose kTWAS and mkTWAS to test if rvTWAS indeed outperform them when analyzing rare variants, which essentially is testing whether SuSiE-based feature selection outperforms the prediction-based feature selection in kTWAS and the marginal test-based feature selection in mkTWAS. Mathematically, the same as rvTWAS, both kTWAS and mkTWAS adopts kernel machine for Step 2 association test: *Q* = *y*^′^*K*_*w*_*y*, where the weighted kernel *K*_*w*_ = *G*^T^*WG*/n. The difference lies in that, in Step 1, kTWAS uses Elastic Net training in standard TWAS for the selection of genetic variants and their weights β_i_: e ∼ Σ β_i_X_i[ref]_ + *ε* (where, the same as the case of PrediXcan, e is the gene expression, X_i[ref]_ is the *i*-th genetic variant in the region, and β_i_ is the coefficient to be estimated); and mkTWAS uses marginal tests of one genetic variant a time to learn the weights β_i_: e ∼ β_i_X_i[ref]_ + *ε*.

### Procedure of Simulations

We conducted simulations to compare rvTWAS to the above methods. Aligning to the rvTWAS protocol borrowing gene expression from a reference dataset, the simulations worked on two datasets: the reference expression dataset and the main GWAS dataset. Based on the GTEx genotype (containing 866 individuals with 51,717,523 SNVs)(GTEx Consortium 2013; GTEx Consortium 2020), we simulated gene expressions in the reference dataset (for rvTWAS Step 1). Based on the 1000 Genomes Project genotype (containing 2,548 subjects with 4,422,985 SNVs) (THE 1000 GENOMES PROJECT CONSORTIUM 2015), we simulated phenotype in the GWAS dataset (for rvTWAS Step 2). In all cases, we simulate a genetic component first and then incorporate it to the expression or phenotypes using prespecified values of genetic component (i.e., heritability).

### Aggregation of the effect of rare variants and its joint effect in the presence of common variants

The aggregation of rare variants is carried out using two alternatives: (1) Following the conventional *heterogeneity* model, which aligns to the hypothesis of “burden test” (Li and Leal 2008), we form “pseudo-SNVs” to represent the aggregated effects of multiple rare variants.

More specifically, a number of (*M* = 0, 50, 100, or 200) rare variants (with MAF <0.5%) in the focal gene region were randomly selected and combined into a pseudo-SNV. The pseudo-SNV is coded by aggregation: the subjects carrying at least one of the *M* rare variants will be coded as 2; and the subjects that do not carry any rare variant be coded as 0. (2) Following an *additive* model (Wu *et al*. 2011), we count the number of rare variants carried by an individual as the contribution of the rare variants to its phenotype, which is equivalent to a “Sum” of these rare variants. Note that homozygous sites will be counted twice.

Acknowledging that common variant may also contribute to gene expression or phenotype, the overall genetic component is modelled as the sum of this rare variant effect (i.e., pseudo-SNV or Sum-effect) and common variants. More specifically, five common variants (with MAF >5%) were randomly selected as causal variants, and the weighted sum of these six causal variants (5 common + 1 pseudo SNVs) will be considered as the genetic component (of the expression or phenotype). The weight of each causal variant is sampled from a standard normal distribution *N*(0,1).

#### Genetic architectures of phenotype and expressions

Two typical genetic architectures are considered: *causality*, where genotypes alter phenotype via the expression, and *pleiotropy*, where genotypes contribute to phenotype and expression independently. Under both scenarios, we simulated expression and phenotypes using an additive genetic model, in which phenotypes and expression are caused by a weighted sum of genetic effects.

We simulate the expressions or phenotypes used the formula below:

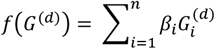

where *G* denotes the genotype matrix, The effect sizes *β*_*i*_ were drawn from the standard normal distribution *N*(0,1) mentioned above. Superscript *d* may be 1 or 2 for reference (GTEx) and GWAS genotype datasets (1000 Genomes Project), respectively. The expression in reference and GWAS datasets thus are generated by:

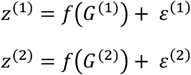

where *ε*^(1)^ and *ε*^(2)^specify the levels of genetic component by adjusting their variance (which will be detailed below).

With a *causality* model, the phenotype is decided by expression:

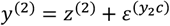

Whereas with a *pleiotropy* model, the phenotype is decided by genetics directly using a similar linear formula for expression (except that the variance component is rescaled by expression heritability instead of trait heritability):

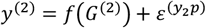

The genetic component contributing to expression or phenotypes is prespecified as a value between 0 and 1 (i.e., heritability). Under both models, the variance components are rescaled by expression heritability and phenotype heritability to ensure the relevant heritability indeed matches the prespecified parameters.

#### Rescaling to match heritability

In the above simulations of expression and phenotypes, the residual *ε* serves as the role to adjust genetic components based on prespecified values (i.e., heritability) denoted as *h*2. To achieve that, we first generated the genetic component based on related formulas and real genotypes, and next calculated its variance of the genetic component as 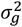. We then solved 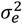 in equation 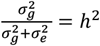. We then sample 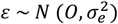 to determine values of residuals, ensuring the heritability to be exactly *h*^2^.

#### Type-I error estimation & power calculations

We generated random phenotypes using for the 1000 Genomes Project individuals (N = 2,548) and analyze them using rvTWAS to generate the empirical null distribution of the p-values. Then the type-I error for the rvTWAS model was estimated using the top 5% cut-off for the most significant p-values.

For each of the genetic architectures and their associated parameters, we generated 1,000 datasets with simulated phenotype and causal variants. The power was then calculated as the proportion of each protocol’s success in identifying the causal gene in each dataset, where the success is defined as a Bonferroni-corrected p-value that is lower than a predetermined critical cut-off of 0.05.

### Real Data Analysis

In addition to the simulations, we also conducted real data analysis to compare alternative methods. The data sources and processing procedures are detailed below.

#### Data source and QC

We run rvTWAS with default parameters on three datasets of brain diseases, schizophrenia (SCZ) (Kiezun *et al*. 2012; Pasaniuc *et al*. 2012), bipolar disorder (BPD) (Kiezun *et al*. 2012; Pasaniuc *et al*. 2012) and autism spectrum disorder (ASD) (Neale *et al*. 2012; DE RUBEIS *et al*. 2014), downloaded from dbGaP web portal (**Data and code availability**). The GTEx (release 8) consortium has provided whole genome sequencing (WGS) and RNA-sequencing (RNA-seq) data for multiple tissues. We included whole blood tissue that has the largest sample size (N=670) in our study. The fully processed, filtered, and normalized gene expression data matrices (in BED format) was downloaded from GTEx portal (**Data and code availability**). The whole genome sequencing file, GTEx_Analysis_2017-06-05_v8_WholeGenomeSeq_866Indiv.vcf was downloaded from dbGaP (**Data and code availability**). Additionally, the following files from dbGaP and GTEx portal are used for quality control: The sample attributes (phs000424.v8.pht002743.v8.p2.c1.GTEx_Sample_Attributes.GRU.txt), the subject phenotypes for sex and age adjustment (phs000424.v8.pht002742.v8.p2.c1.GTEx_Subject_Phenotypes.GRU.txt), the covariates adjustments in eQTL analysis, including genotyping principal components (PCs) (GTEx_Analysis_v8_eQTL_covariates.tar.gz). Based on the GTEx recommendation for whole blood sample, 60 PEER factors (Stegle *et al*. 2012) were regressed out for adjustment of additional confounding factors.

The genotype QC was conducted by PLINK (Purcell *et al*. 2007). The QC steps involve excluding variants with missingness rate > 0.1, high deviations from Hardy-Weinberg equilibrium (P-value >10^−6^) and removing samples with missingness rate > 0.1. For genotype datasets containing only the common variants with minor allele frequency (MAF)>0.05, we used “--maf 0.05” to filter out all variants with MAF below 0.05, while for genotype datasets containing only the rare variants with MAF < 0.005, we used “--max-maf 0.005” to impose an upper MAF bound as 0.005. Eventually, we collected 11,214 individuals with 1,580,125 SNVs for SCZ, 7,411 individuals with 1,581,749 SNVs for BPD, and 7,766 individuals with 3,158,065 SNVs for ASD.

When aligning two datasets’ genotype coordinates, i.e., the reference dataset (GTEx) and GWAS dataset (the three disorder cohorts), Liftover (Hinrichs *et al*. 2006) was utilized to convert all coordinates to HG38. The locations that are in overlap were used for the analyses.

#### Functional validations of identified genes

To search the evidence whether the identified genes are related to the disease’s susceptibility, we performed gene-disease association analysis using the DisGeNET repository (v7.0) (Pinero *et al*. 2015; Pinero *et al*. 2017; Pinero *et al*. 2020), a platform containing 1,134,942 gene-disease associations (GDAs) between 21,671 genes and 30,170 traits. For each disease, DisGeNET has its specific summary of gene-disease associations containing information of identified significant genes and their GAD scores. We assessed each method under comparison based on the number of its discovered genes which are reported as disease associated in DisGeNET (*validated*), as well as the proportion (*validated rate*) of these validated genes among all the genes identified (i.e., validated genes divided by total number of identified genes). We also examine the number and proportion of uniquely discovered genes which have annotations. Specifically, for any pair of methods A and B yielding corresponding sets of associated genes, we take the difference of the sets A - B (genes only in A and not B), and B - A (genes only in B but not A), and find the number of genes (*uniquely validated*) as well as the proportion of genes (*uniquely validated rate*) in each set difference which are annotated in DisGeNET.

## RESULTS

### Simulations show rvTWAS has a well-controlled type-I error and higher power

Our null simulation shows that the type-I error of rvTWAS, i.e., the 5% cut-off (determined by simulating traits under the null distribution) is 0.051, which is close to the targeted type-I error of α = 0.05. Therefore, the type I error of rvTWAS is under control. The type-I-errors of kTWAS and mkTWAS have been shown previously (Cao *et al*. 2021b; Cao *et al*. 2022).

With the type-I error controlled, we then compare power. Under pleiotropy, rvTWAS significantly outperform PrediXcan that does not disentangle feature selection and aggregation, as well as kTWAS and mkTWAS that do not use Bayesian feature selections, in both heterogeneity (*pseudo-SNV*) and additive models (*Sum*) (**Figure 2; Supplementary Figure 1**). When there is no causal rare variant, rvTWAS is slightly more powerful than the alternatives (**Figure 2A; Supplementary Figure 1A**), while the advantage is more pronounced when more rare variants are included as causal variants (**Figure 2B,C,D**; **Supplementary Figure 1B**,**C**,**D**). All kernel-based methods (rvTWAS, kTWAS and mkTWAS) outperform the GReX-based protocols (PrediXcan), evidencing the supremacy of kernel-based methods in feature aggregation. Under causality scenario, the observation is similar to the pleiotropy model: rvTWAS outperforms other tools and the effect is more pronounced when more rare variants are present (**Figure 3; Supplementary Figure 2**). All models enjoy an observable higher power in pleiotropy than causality, which is consistent with previous simulations (Veturi AND Ritchie 2018; Cao *et al*. 2021a; Tang *et al*. 2021).

**Figure 2.**
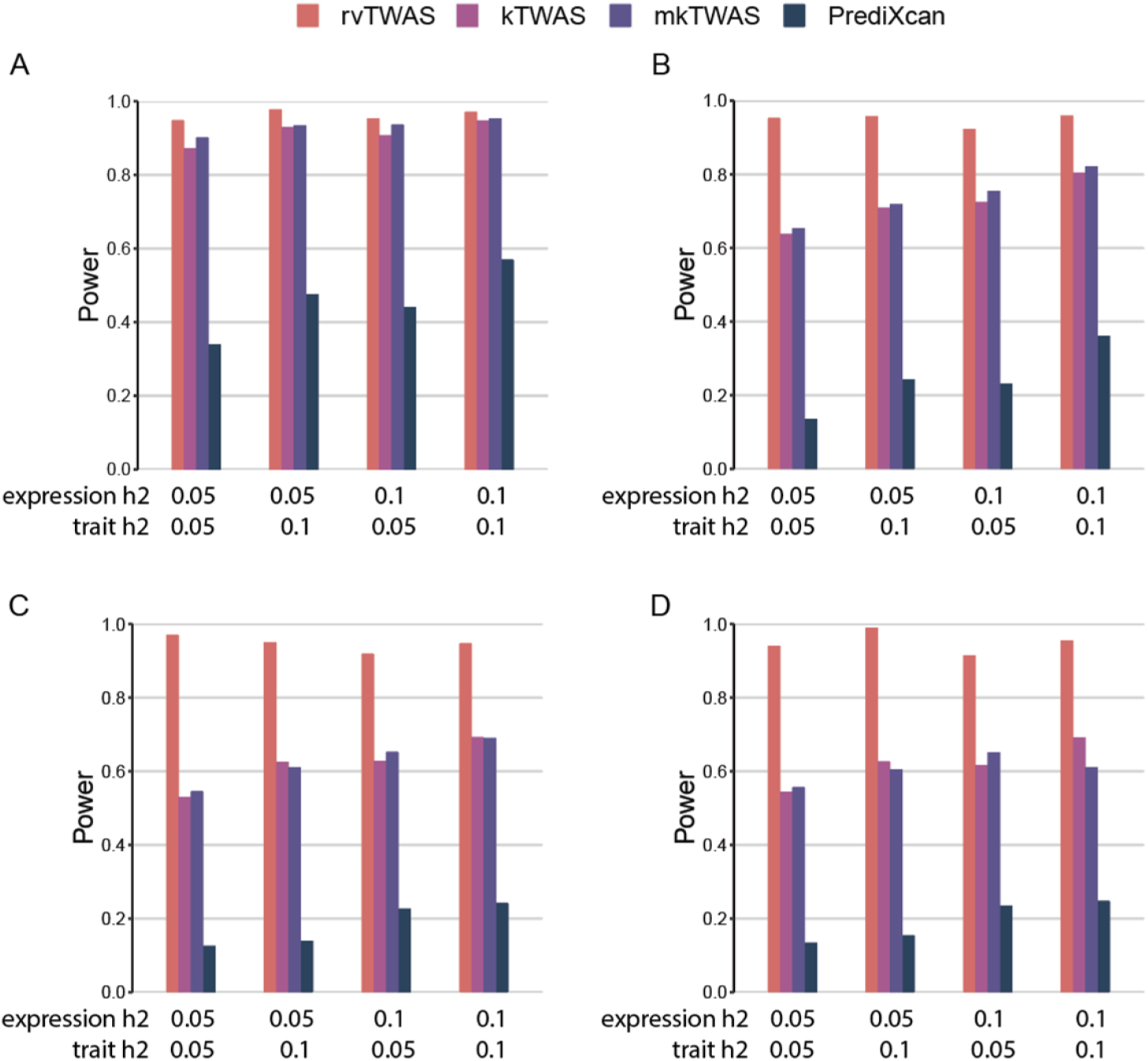
Power comparison under pleiotropy scenarios. Power is indicated on the y-axis. All panels are results under an additive genetic architecture, with differing expression heritability and trait heritability denoted below each panel. The total number of contributing rare genetic variants (with MAF < 0.005) M are fixed in each panel. **A**) M=0; **B**) M=50; **C**) M=100; and **D**) M=200.

**Figure 3.**
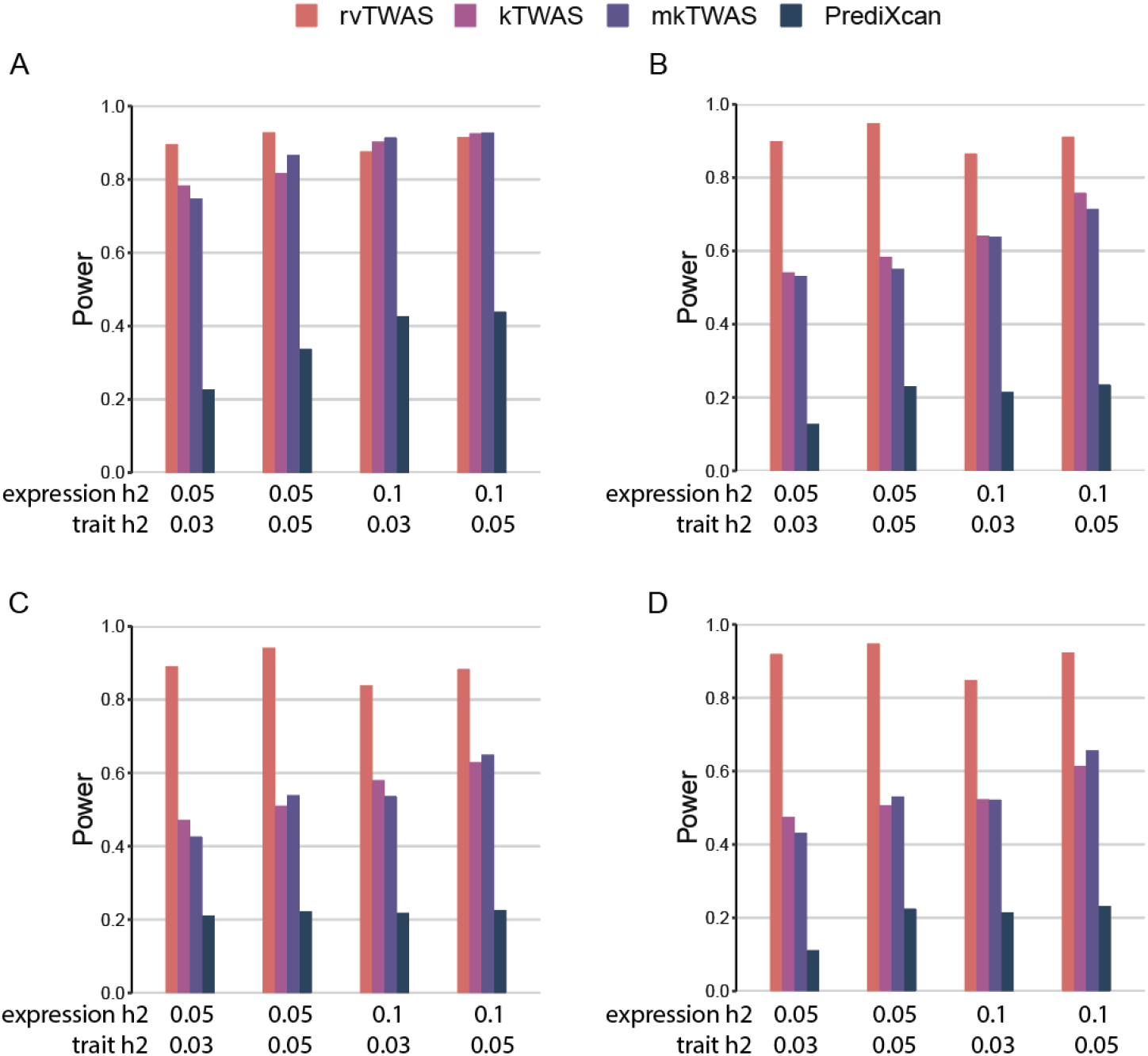
Power comparison under causality scenarios. Power is indicated on the y-axis. All panels are results under an additive genetic architecture, with differing expression heritability and trait heritability denoted below each panel. The total number of contributing rare genetic variants (with MAF < 0.005) M are fixed in each panel. **A**) M=0; **B**) M=50; **C**) M=100, and **D**) M=200.

### rvTWAS outperforms alternative methods in real data analysis of SCZ, BPD, and ASD

The significant genes identified by the four methods to three brain disorders are presented in **Supplementary Tables 1-3**. Below is the analysis of their indications.

### rvTWAS identifies more genes

The Manhattan plots generated by applying rvTWAS are shown in **Figure 4**, and the corresponding Manhattan plots generated by alternative methods are in **Supplementary Figures 3-5**. Evidently, rvTWAS outperforms alternative in terms of the number of genes identified. In SCZ, at a Bonferroni-corrected P < 0.05, rvTWAS identified 30 susceptibility genes, which is more than those identified by PrediXcan (7 genes), kTWAS (24 genes) and mkTWAS (22 genes). We conducted similar comparisons for BPD and ASD, and demonstrated consistent trends of more genes identified by rvTWAS compared to other approaches. For BPD, rvTWAS identified 127 susceptibility genes whereas 53, 8, and 64 genes were identified by PrediXcan, kTWAS and mkTWAS, respectively. For ASD, rvTWAS identified 41 susceptibility genes while 41, 19, and 29 genes were identified for PrediXcan, kTWAS and mkTWAS, respectively (**Supplementary Figure 6**).

**Figure 4.**
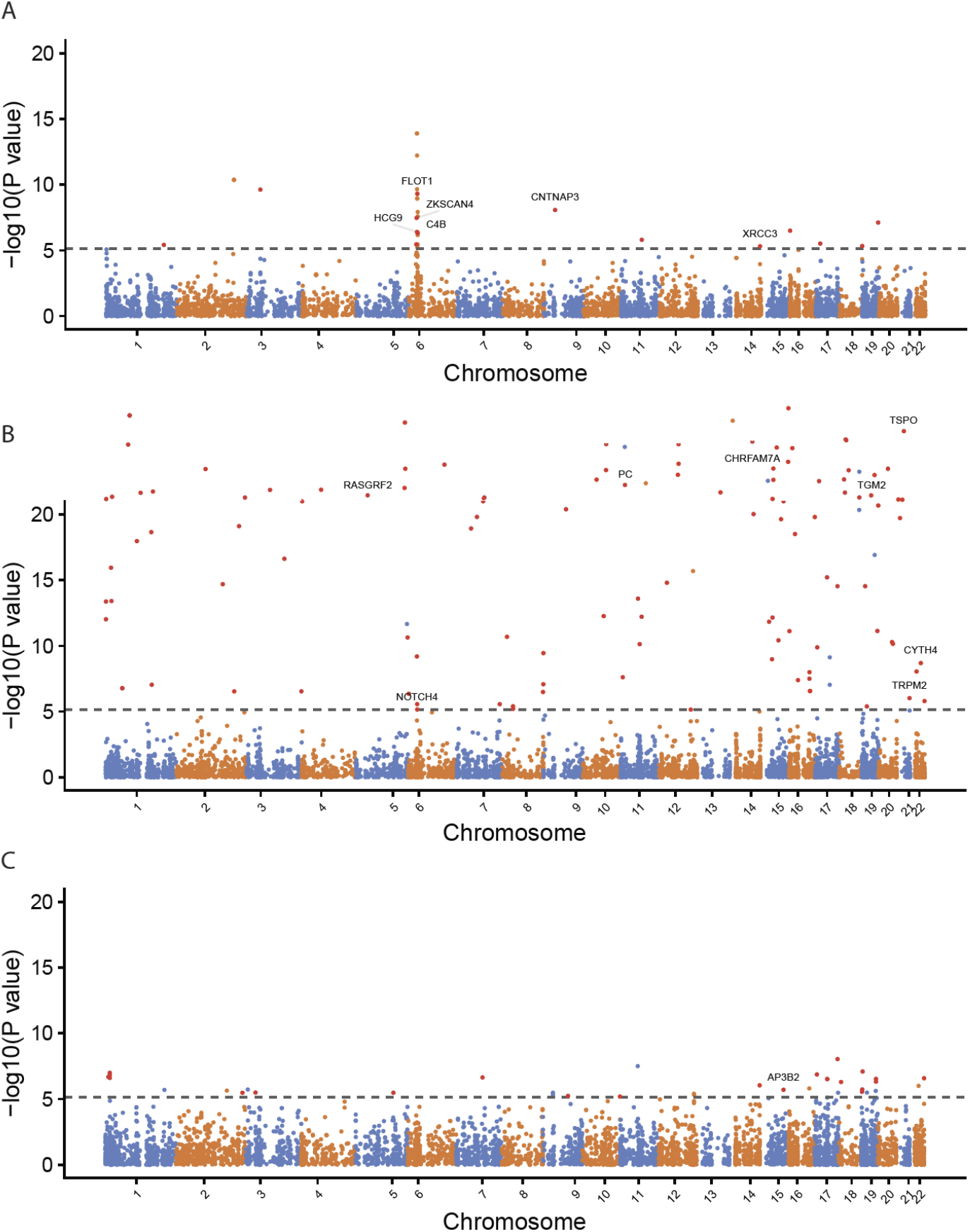
Manhattan plots showing associations identified by rvTWAS. Black dashed lines indicate the cut-off specified by Bonferroni correction at the level of P-value < 0.05. Red dots represent uniquely identified susceptibility genes. The uniquely identified signals that have been reported literatures are labeled by its gene name. The Y-axis ranges from 0 to 20 for a convenient visualization. The signals with -log10(P-value) larger than 20 can be found in **Supplementary Tables 1-3. A)** SCZ. **B)** BPD. **C)** ASD.

### Genes uniquely identified by rvTWAS are supported by literatures

In SCZ, among the 30 genes, 14 genes were uniquely identified by rvTWAS. Out of these 14 uniquely identified, 6 have been reported to be associated with SCZ; they are ZKSCAN4 (nominal P=3.37e-08) (Yue *et al*. 2011), HCG9 (nominal P=3.94e-07) (Pal *et al*. 2016), FLOT1 (nominal P=4.88e-10) (Buxton *et al*. 2019; Zhong *et al*. 2019), C4B (nominal P=4.79e-07) (Mayilyan *et al*. 2008; Sekar *et al*. 2016), CNTNAP (nominal P=3.84e-09) (Ji *et al*. 2013) and XRCC3 (nominal P=4.80e-06) (Odemis *et al*. 2016). Similar outcomes are observed for BPD and ASD. Eight uniquely identified genes in BPD are supported by literatures, including RASGRF2 (nominal P=2.65e-24) (Stacey *et al*. 2012; Douglas *et al*. 2016), NOTCH4 (nominal P=6.98e-06) (Dieset *et al*. 2012), PC (nominal P=1.81e-32) (Knowles *et al*. 2017), CHRFAM7A (nominal P=3.46e-181) (Flomen *et al*. 2006; Kunii *et al*. 2015), TGM2 (nominal P=3.17e-24) (Graae *et al*. 2012; Gupta *et al*. 2018; Vastrad AND Vastrad 2022), TRPM2 (nominal P=9.54e-07) (Jang *et al*. 2015), CYTH4 (nominal P=2.05e-09) (Rezazadeh *et al*. 2015), and TSPO (nominal P=3.44e-211) (Scaini *et al*. 2019), and one uniquely identified gene in ASD is supported by literature, AP3B2 (nominal P=1.98e-06) (O’ROAK *et al*. 2012).

### rvTWAS identified genes have higher rates of validation in DisGeNET

To quantitively assess the functional relevance of the identified genes, we referred to the DisGeNET database of human gene-disease associations (Pinero *et al*. 2015; Pinero *et al*. 2017; Pinero *et al*. 2020). We assessed each protocol on the number and proportion of its identified genes that are reported as disease associated in DisGeNET (*validated* and *validated rate*), as well as the corresponding statistics for uniquely identified (**Materials and Methods**). We found that rvTWAS outperforms all alternatives for SCZ and BPD with a very large margin, although slightly lower than PrediXcan in ASD by 2 vs. 3 (**Figure 5**). These results suggest that rvTWAS is powerful in uncovering additional susceptibility genes that might have been missed out by other approaches.

**Figure 5:**
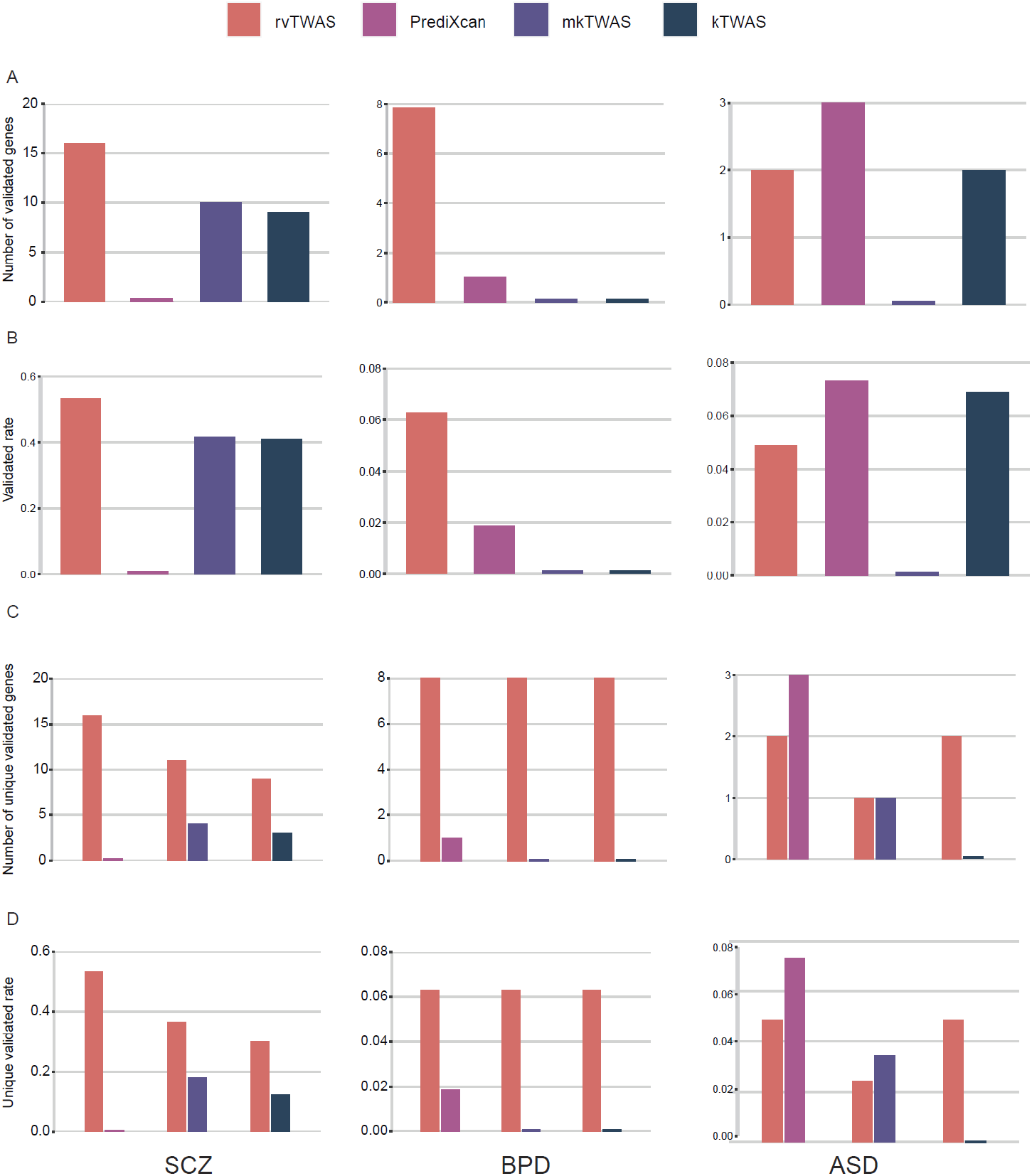
The number and proportion of genes identified by rvTWAS and other methods that are validated by DisGeNET. **A)** Number of validated genes. **B)** Proportion of validated genes (validated rate). **C)** Number of validated genes reported exclusively by one of the two protocols in the DisGeNET database. **D)** Proportion (unique validated rate) of genes reported exclusively by one of the two protocols in the DisGeNET database. For all panels, the left, middle, and right columns are for SCZ, BPD and ASD, respectively.

## DISCUSSION

The outcome of rvTWAS presented in **Results** are based on inclusive analysis of rare variants and common variants. So, the effect is not only caused by rare variants. One may be curious about the contribution of rare variants to the overall results, as well as the performance of rvTWAS on rare variants only analysis.

To understand the contribution of different spectrums of data, for the significant genes identified by rvTWAS, we specifically looked at the genetic variants selected by SuSiE. Interestingly, for BPD most genes (70%) contain rare variants (**Figure 6A** Middle Column), whereas for ASD most genes (88%) only contain common variants (**Figure 6A** Right Column). The proportion SCZ is in between with a moderate percentage (33%) containing rare variants (**Figure 6A** Left Column).

**Figure 6.**
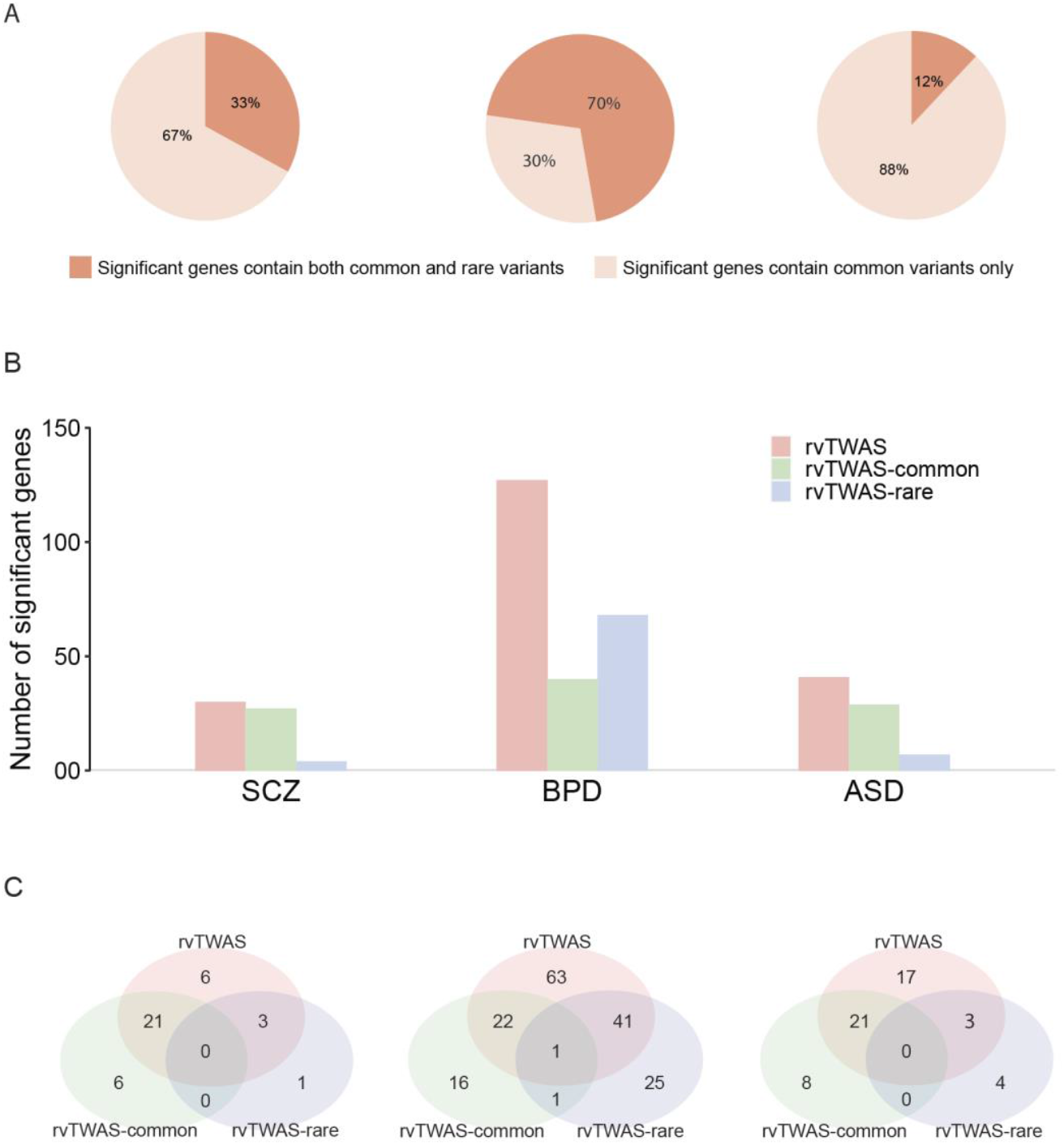
Contributions of rare and common variants to the outcome of rvTWAS. **A**. The percentage of significant genes for which SuSiE have selected both rare and common variants *vs*. genes for which only common variants have been selected. **B**. The number of significant genes discovered by rvTWAS, rvTWAS-common and rvTWAS-rare. **C**. The numbers of overlapped genes among rvTWAS (including both common and rare variants) and the two filtered rvTWAS analyses, rvTWAS-common and rvTWAS-rare. For all panels, the left, middle, and right columns are for SCZ, BPD and ASD, respectively.

To understand what will happen if the common and rare variants are separately analyzed, we specifically carried out two additional analyses: 1) by specifying a minimal MAF cut-off of 0.05, only common genetic variants are included, which is termed as rvTWAS-common; 2) by specifying a maximal MAF cut-off of 0.005, only rare variants are included, which is called rvTWAS-rare. These additional protocols are compared to the main model: 3) the inclusive protocol without any filters, which is just called rvTWAS. Note that the filtering of data was conducted before the SuSiE-based Bayesian feature selection (**Materials & Methods**). There are two observations. First, rvTWAS evidently identified more genes than either rvTWAS-common or rvTWAS-rare alone in all three disorders. Moreover, rvTWAS identified number is higher than the aggregated numbers of rvTWAS-common *and* rvTWAS-rare in both ASD and BPD (**Figure 6B**). This suggests that modeling the joint contribution of common and rare variants outperforms splitting them. Second, while the overlap between rvTWAS-common and rvTWAS-rare are universally low, the overlap between rvTWAS and an alternative is depending on specifical disease (**Figure 6C**). Interestingly, in BPD, a significant number of rare-variant only genes are identified, however this is not the case for SCZ and ASD. Analysis of the contributions of rare and common variants to the psychiatric disorder have been attempted by others (Reay AND Cairns 2020; SUL *et al*. 2020; Dattani *et al*. 2023; Mollon *et al*. 2023). Using rvTWAS, more thorough characterization may be carried out in the future work.

Playing a critical role in Step 2 of rvTWAS, SKAT (Wu *et al*. 2011; Lee *et al*. 2012) is a flagship tool for rare variants analysis using kernel machine, although it does not utilize gene expressions. In practice, when running SKAT to rare variant analysis, one often needs some criteria of filtering, otherwise there will be too many results. For instance, using SKAT to analyze the three datasets of psychiatric disorders without any filtering yields very large numbers of identified genes (with 916 for SCZ, 4419 for BPD and 851 for ASD), which look untrue. Therefore, rvTWAS provides a transcriptome-directed strategy in prioritizing and weighting rare variants for SKAT.

A limitation of rvTWAS lies is that the rare variants cannot be too rare (e.g., singletons private to the carrying individual). As the cohorts of feature selection (such as GTEx) and association mapping are different, one relies on their sharing the same locations or an imputation (Das *et al*. 2016). The very rare variants will have poor sharing, which are also difficult to impute. The spectrum of frequencies that allows optimal use of rvTWAS may be characterized in the future.

In conclusion, we developed rvTWAS, a rare-variant-focused tool leveraging our unique insight into disentangling TWAS protocols into feature selection and aggregation. Through simulations and real data analysis, we showed that rvTWAS outperforms alternative protocols in the TWAS domain. Future work includes the extension of rvTWAS to the use of other middle traits such as brain imaging (Xu *et al*. 2017). Methods of feature aggregation other than kernel may also be tested.

## Author contributions

Conceived the idea and designed the models: Q.Z. Analyzed real data: J.H., Q.L., and Q.Z. Developed the software: J.H. Conducted simulations: J.H. Wrote the paper: J.H. and Q.Z. Supervised the project: Q.Z.

## Data and code availability

rvTWAS, https://github.com/theLongLab/rvTWAS

mkTWAS, https://github.com/theLongLab/mkTWAS

kTWAS, https://github.com/theLongLab/kTWAS

PrediXcan, https://github.com/hakyim/PrediXcan

SuSiE, https://github.com/stephenslab/susieR

PLINK, https://zzz.bwh.harvard.edu/plink/

1000 Genomes Project, https://www.internationalgenome.org/

GTEx, https://gtexportal.org/ and https://www.ncbi.nlm.nih.gov/projects/gap/cgi-bin/study.cgi?study_id=phs000424.v8.p2

Exon sequencing data of Schizophrenia, https://www.ncbi.nlm.nih.gov/projects/gap/cgi-bin/study.cgi?study_id=phs000473.v2.p2

Autism Sequencing Consortium (ASC), https://www.ncbi.nlm.nih.gov/projects/gap/cgi-bin/study.cgi?study_id=phs000298.v4.p3

Exon sequencing data of bipolar disorder, https://www.ncbi.nlm.nih.gov/projects/gap/cgi-bin/study.cgi?study_id=phs000473.v2.p2

## Finding

Q.Z. is supported by an NSERC Discovery Grant (RGPIN-2018-05147), a University of Calgary VPR Catalyst grant, and a New Frontiers in Research Fund (NFRFE-2018-00748). J.H. is partly supported by an Alberta Innovates LevMax-Health Program Bridge Funds (222300769) and a CSC scholarship. The computational infrastructure is funded by a Canada Foundation for Innovation JELF grant (36605) and an NSERC RTI grant (RTI-2021-00675).

