## Supplementary materials for "rvTWAS: identifying gene-trait association using sequences by utilizing transcriptome-directed feature selection"

### Supplementary Tables

**Supplementary Table 1. Susceptibility genes identified by rvTWAS, PrediXcan, kTWAS and mkTWAS, at a Bonferroni-corrected P < 0.05 for Schizophrenia (SCZ)**

| Model | Gene | Chr | Start | End | P-value | Validated |
|---|---|---|---|---|---|---|
| rvTWAS | PPP1R12B | 1 | 202348699 | 202590595 | 3.77E-06 | - |
| rvTWAS | FTCDNL1 | 2 | 199760544 | 199851173 | 4.25E-11 | + |
| rvTWAS | C2orf69 | 2 | 199911256 | 199955392 | 4.25E-11 | - |
| rvTWAS | MAIP1 | 2 | 199955317 | 200008540 | 4.25E-11 | + |
| rvTWAS | ZMYND10 | 3 | 50341229 | 50346852 | 2.36E-10 | - |
| rvTWAS | ZNF391 | 6 | 27374615 | 27403904 | 3.47E-06 | - |
| rvTWAS | ZKSCAN4 | 6 | 28244623 | 28252224 | 3.37E-08 | + |
| rvTWAS | GABBR1 | 6 | 29602228 | 29633976 | 1.26E-14 | + |
| rvTWAS | ZFP57 | 6 | 29672392 | 29681110 | 1.27E-14 | - |
| rvTWAS | HLA-F-AS1 | 6 | 29735028 | 29748993 | 2.21E-10 | - |
| rvTWAS | HLA-G | 6 | 29826967 | 29831125 | 6.01E-13 | + |
| rvTWAS | HCP5B | 6 | 29871895 | 29873783 | 1.11E-09 | - |
| rvTWAS | HCG9 | 6 | 29975112 | 29978410 | 3.94E-07 | + |
| rvTWAS | FLOT1 | 6 | 30727709 | 30742733 | 4.88E-10 | + |
| rvTWAS | IER3 | 6 | 30743199 | 30744554 | 5.05E-10 | - |
| rvTWAS | LINC00243 | 6 | 30798654 | 30830659 | 1.15E-09 | + |
| rvTWAS | VARS2 | 6 | 30914205 | 30926459 | 6.10E-06 | + |
| rvTWAS | C4A | 6 | 31982024 | 32002681 | 4.49E-07 | + |
| rvTWAS | C4B | 6 | 32014762 | 32035418 | 4.79E-07 | + |
| rvTWAS | CYP21A2 | 6 | 32038265 | 32041670 | 6.74E-07 | + |
| rvTWAS | PPT2 | 6 | 32153999 | 32163680 | 3.53E-06 | - |
| rvTWAS | AGER | 6 | 32180968 | 32184324 | 1.19E-08 | + |
| rvTWAS | PBX2 | 6 | 32184836 | 32190186 | 2.57E-08 | + |
| rvTWAS | CNTNAP3 | 9 | 39072767 | 39288315 | 8.40E-09 | + |
| rvTWAS | KRTAP5-9 | 11 | 71548418 | 71549553 | 1.54E-06 | - |
| rvTWAS | XRCC3 | 14 | 103697609 | 103715504 | 4.80E-06 | + |
| rvTWAS | WDR90 | 16 | 649325 | 667833 | 3.16E-07 | - |
| rvTWAS | ZSWIM7 | 17 | 15976560 | 15993830 | 3.02E-06 | - |
| rvTWAS | HSBP1L1 | 18 | 79964561 | 79970810 | 4.63E-06 | - |
| rvTWAS | FIZ1 | 19 | 55591371 | 55601878 | 7.57E-08 | - |
| PrediXcan | SCAP | 3 | 47413694 | 47477126 | 5.18E-08 | - |
| PrediXcan | ZNF699 | 19 | 9294275 | 9309838 | 1.86E-07 | - |
| PrediXcan | PP7080 | 5 | 466124 | 473098 | 2.16E-07 | - |
| PrediXcan | PDLIM3 | 4 | 185500660 | 185535612 | 5.00E-07 | - |
| PrediXcan | TRIM35 | 8 | 27284887 | 27311319 | 8.20E-07 | - |
| PrediXcan | SPECC1L | 22 | 24270817 | 24417740 | 3.91E-06 | - |
| PrediXcan | VN1R1 | 19 | 57454790 | 57457142 | 4.02E-06 | - |

| | | | | | | |
|---|---|---|---|---|---|---|
| kTWAS | SLC35E2B | 1 | 1659529 | 1692728 | 1.57E-08 | - |
| kTWAS | SLC35E2 | 1 | 1724838 | 1745992 | 1.88E-07 | - |
| kTWAS | SYTL1 | 1 | 27342020 | 27353932 | 3.37E-08 | - |
| kTWAS | EML6 | 2 | 54723665 | 54972025 | 1.09E-06 | - |
| kTWAS | CLHC1 | 2 | 55174791 | 55232354 | 4.58E-06 | - |
| kTWAS | TMEM163 | 2 | 134455759 | 134719000 | 5.45E-06 | - |
| kTWAS | MAIP1 | 2 | 199955317 | 200008540 | 4.25E-11 | + |
| kTWAS | C4orf33 | 4 | 129093608 | 129116640 | 4.75E-06 | - |
| kTWAS | BTN3A2 | 6 | 26365159 | 26378320 | 2.19E-07 | + |
| kTWAS | GABBR1 | 6 | 29602228 | 29633976 | 1.26E-14 | + |
| kTWAS | ZFP57 | 6 | 29672392 | 29681110 | 1.26E-14 | - |
| kTWAS | HLA-G | 6 | 29826967 | 29831125 | 1.62E-07 | + |
| kTWAS | HCP5B | 6 | 29871895 | 29873783 | 2.19E-06 | - |
| kTWAS | IER3 | 6 | 30743199 | 30744554 | 4.69E-10 | - |
| kTWAS | VARS2 | 6 | 30914205 | 30926459 | 6.08E-06 | + |
| kTWAS | CCHCR1 | 6 | 31142439 | 31158238 | 1.26E-08 | + |
| kTWAS | POU5F1 | 6 | 31164337 | 31180731 | 4.28E-06 | + |
| kTWAS | CLIC1 | 6 | 31730618 | 31739763 | 5.12E-10 | - |
| kTWAS | C4A | 6 | 31982024 | 32002681 | 1.58E-07 | + |
| kTWAS | PPT2 | 6 | 32153999 | 32163680 | 4.69E-06 | - |
| kTWAS | AGER | 6 | 32180968 | 32184324 | 2.57E-08 | + |
| kTWAS | PBX2 | 6 | 32184836 | 32190186 | 5.56E-08 | + |
| kTWAS | RP11-890B15.3 | 11 | 130866254 | 130870247 | 3.15E-06 | - |
| kTWAS | PPP2R5C | 14 | 101761798 | 101927989 | 1.70E-06 | - |
| mkTWAS | FTCDNL1 | 2 | 199760544 | 199851173 | 4.25E-11 | + |
| mkTWAS | C2orf69 | 2 | 199911256 | 199955392 | 3.66E-10 | - |
| mkTWAS | ZSCAN12 | 6 | 28378955 | 28399734 | 1.78E-09 | + |
| mkTWAS | OR2H2 | 6 | 29587455 | 29589038 | 2.37E-12 | - |
| mkTWAS | MOG | 6 | 29656981 | 29672372 | 1.83E-14 | + |
| mkTWAS | ZFP57 | 6 | 29672392 | 29681110 | 3.77E-13 | - |
| mkTWAS | HLA-F-AS1 | 6 | 29735028 | 29748993 | 1.22E-08 | - |
| mkTWAS | HLA-G | 6 | 29826967 | 29831125 | 1.31E-12 | + |
| mkTWAS | HCP5B | 6 | 29871895 | 29873783 | 2.16E-15 | - |
| mkTWAS | PRR3 | 6 | 30557175 | 30563723 | 4.12E-13 | - |
| mkTWAS | IER3 | 6 | 30743199 | 30744554 | 2.78E-07 | - |
| mkTWAS | HCG20 | 6 | 30766825 | 30792250 | 1.34E-14 | - |
| mkTWAS | LINC00243 | 6 | 30798654 | 30830659 | 3.03E-12 | + |
| mkTWAS | CCHCR1 | 6 | 31142439 | 31158238 | 1.15E-09 | + |
| mkTWAS | MIR6891 | 6 | 31355224 | 31355316 | 2.43E-06 | - |
| mkTWAS | APOM | 6 | 31655471 | 31658210 | 4.15E-09 | - |
| mkTWAS | C4A | 6 | 31982024 | 32002681 | 1.89E-10 | + |
| mkTWAS | CYP21A2 | 6 | 32038265 | 32041670 | 5.90E-11 | + |
| mkTWAS | FKBPL | 6 | 32128707 | 32130291 | 6.98E-08 | - |

| | | | | | | |
|---|---|---|---|---|---|---|
| mkTWAS | NOTCH4 | 6 | 32194843 | 32224067 | 1.49E-06 | + |
| mkTWAS | LINC00239 | 14 | 101730437 | 101732522 | 1.62E-06 | - |
| mkTWAS | SIRPB2 | 20 | 1470741 | 1491587 | 2.47E-07 | - |

**Supplementary Table 2. Susceptibility genes identified by rvTWAS, PrediXcan, kTWAS and mkTWAS, at a Bonferroni-corrected P < 0.05 for Bipolar Disorder (BPD)**

| Model | Gene | Chr | Start | End | P-value | Validated |
|---|---|---|---|---|---|---|
| rvTWAS | DVL1 | 1 | 1335307 | 1349350 | 9.51E-13 | - |
| rvTWAS | ATAD3B | 1 | 1471769 | 1496202 | 4.24E-14 | - |
| rvTWAS | UBR4 | 1 | 19076740 | 19210276 | 1.14E-16 | - |
| rvTWAS | PINK1-AS | 1 | 20642657 | 20651766 | 3.89E-14 | - |
| rvTWAS | SERINC2 | 1 | 31409565 | 31434680 | 2.35E-21 | - |
| rvTWAS | PRPF38A | 1 | 52404564 | 52420839 | 4.06E-23 | - |
| rvTWAS | OMA1 | 1 | 58415384 | 58546802 | 1.64E-07 | - |
| rvTWAS | TACSTD2 | 1 | 58575423 | 58577773 | 1.72E-07 | - |
| rvTWAS | RP11-63G10.4 | 1 | 58715609 | 58771295 | 1.72E-07 | - |
| rvTWAS | CELSR2 | 1 | 109250019 | 109275750 | 1.57E-200 | - |
| rvTWAS | GSTM4 | 1 | 109656081 | 109665496 | 1.08E-18 | - |
| rvTWAS | TRIM33 | 1 | 114392777 | 114511160 | 1.16E-223 | - |
| rvTWAS | BCAS2 | 1 | 114567557 | 114581639 | 1.16E-223 | - |
| rvTWAS | S100A4 | 1 | 153543613 | 153550136 | 1.27E-76 | - |
| rvTWAS | RP11-226L15.5 | 1 | 160024953 | 160026794 | 2.24E-19 | - |
| rvTWAS | FCGR3B | 1 | 161623196 | 161631963 | 9.14E-08 | - |
| rvTWAS | ZBTB41 | 1 | 197153680 | 197200542 | 1.30E-77 | - |
| rvTWAS | DARS | 2 | 135906677 | 135985877 | 3.81E-45 | - |
| rvTWAS | KCNH7 | 2 | 162371596 | 162838730 | 2.04E-15 | - |
| rvTWAS | BMPR2 | 2 | 202376936 | 202567751 | 2.92E-07 | - |
| rvTWAS | CAND2 | 3 | 12796472 | 12871916 | 4.06E-50 | - |
| rvTWAS | PDCD6IP | 3 | 33798352 | 33869707 | 1.65E-22 | - |
| rvTWAS | PARP14 | 3 | 122680618 | 122730840 | 2.14E-122 | - |
| rvTWAS | ATP2C1 | 3 | 130850595 | 131016712 | 1.66E-118 | - |
| rvTWAS | PCCB | 3 | 136250306 | 136336232 | 2.37E-17 | - |
| rvTWAS | MUC20 | 3 | 195720882 | 195741123 | 2.90E-07 | - |
| rvTWAS | PCGF3 | 4 | 705748 | 770640 | 0 | - |
| rvTWAS | TBCK | 4 | 106041599 | 106321495 | 9.83E-29 | - |
| rvTWAS | RASGRF2 | 5 | 80960672 | 81230156 | 2.65E-24 | + |
| rvTWAS | FAM153A | 5 | 177707981 | 177783398 | 2.19E-12 | - |
| rvTWAS | MRNIP | 5 | 179835133 | 179858887 | 2.31E-11 | - |
| rvTWAS | SERPINB1 | 6 | 2832332 | 2842006 | 4.49E-07 | - |
| rvTWAS | MDC1 | 6 | 30699807 | 30717889 | 4.15E-30 | - |
| rvTWAS | POU5F1 | 6 | 31164337 | 31180731 | 6.45E-10 | - |
| rvTWAS | SAPCD1 | 6 | 31762835 | 31764851 | 5.01E-218 | - |
| rvTWAS | CYP21A2 | 6 | 32038265 | 32041670 | 2.73E-06 | - |
| rvTWAS | NOTCH4 | 6 | 32194843 | 32224067 | 6.98E-06 | + |
| rvTWAS | RPS18 | 6 | 33272048 | 33276510 | 2.25E-45 | - |

| | | | | | | |
|---|---|---|---|---|---|---|
| rvTWAS | INTS1 | 7 | 1470277 | 1504367 | 1.81E-142 | - |
| rvTWAS | IKZF1 | 7 | 50304124 | 50405101 | 1.18E-19 | - |
| rvTWAS | RBM48 | 7 | 92528773 | 92538005 | 0 | - |
| rvTWAS | CAPZA2 | 7 | 116811070 | 116922049 | 1.97E-57 | - |
| rvTWAS | TRBC1 | 7 | 142791694 | 142793368 | 2.13E-22 | - |
| rvTWAS | TRBC2 | 7 | 142801041 | 142802748 | 1.71E-22 | - |
| rvTWAS | GIMAP6 | 7 | 150625375 | 150632648 | 2.75E-06 | - |
| rvTWAS | CNOT7 | 8 | 17224964 | 17246570 | 2.05E-11 | - |
| rvTWAS | BAG4 | 8 | 38176731 | 38213301 | 3.98E-06 | - |
| rvTWAS | RP11-350N15.5 | 8 | 38382364 | 38383461 | 6.17E-06 | - |
| rvTWAS | TOP1MT | 8 | 143304384 | 143359979 | 3.24E-07 | - |
| rvTWAS | FBXL6 | 8 | 144355431 | 144358512 | 8.35E-08 | - |
| rvTWAS | ZNF251 | 8 | 144720907 | 144755605 | 3.58E-10 | - |
| rvTWAS | MIGA2 | 9 | 129036621 | 129072082 | 1.55E-63 | - |
| rvTWAS | MRPS16 | 10 | 73248863 | 73252693 | 5.49E-13 | - |
| rvTWAS | COX15 | 10 | 99711844 | 99732100 | 9.57E-37 | - |
| rvTWAS | ADAM8 | 10 | 133262403 | 133276868 | 3.87E-138 | - |
| rvTWAS | SIGIRR | 11 | 405716 | 417397 | 2.98E-115 | - |
| rvTWAS | DNHD1 | 11 | 6497296 | 6572022 | 2.44E-08 | - |
| rvTWAS | MS4A3 | 11 | 60056587 | 60071128 | 2.56E-14 | - |
| rvTWAS | EFEMP2 | 11 | 65866441 | 65873592 | 7.48E-11 | - |
| rvTWAS | RIN1 | 11 | 66332062 | 66336840 | 1.15E-198 | - |
| rvTWAS | PC | 11 | 66848233 | 66958376 | 1.81E-32 | + |
| rvTWAS | ARAP1 | 11 | 72685069 | 72752403 | 6.16E-13 | - |
| rvTWAS | ZNF384 | 12 | 6666477 | 6689510 | 3.43E-84 | - |
| rvTWAS | TM7SF3 | 12 | 26973195 | 27014434 | 1.59E-15 | - |
| rvTWAS | RP3-424M6.4 | 12 | 110501614 | 110503441 | 7.03E-06 | - |
| rvTWAS | RP11-131L12.3 | 12 | 118428281 | 118428870 | 2.04E-16 | - |
| rvTWAS | COQ5 | 12 | 120503274 | 120534350 | 7.14E-91 | - |
| rvTWAS | ABCB9 | 12 | 122920951 | 122981570 | 1.19E-99 | - |
| rvTWAS | ARL6IP4 | 12 | 122981650 | 122982913 | 3.36E-115 | - |
| rvTWAS | DHRS4 | 14 | 23955035 | 23969279 | 1.97E-120 | - |
| rvTWAS | ATP6V1D | 14 | 67294371 | 67359996 | 1.69E-219 | - |
| rvTWAS | CHRFAM7A | 15 | 30357766 | 30393849 | 3.46E-181 | + |
| rvTWAS | GOLGA8R | 15 | 30403740 | 30414162 | 1.47E-12 | - |
| rvTWAS | GOLGA8B | 15 | 34525207 | 34583651 | 1.06E-59 | - |
| rvTWAS | C15orf62 | 15 | 40770290 | 40772449 | 1.06E-09 | - |
| rvTWAS | CAPN3 | 15 | 42359500 | 42412317 | 7.09E-13 | - |
| rvTWAS | RPS27L | 15 | 63125872 | 63158021 | 3.79E-11 | - |
| rvTWAS | RP11-775C24.5 | 15 | 80999593 | 80999981 | 0 | - |

| | | | | | | |
|---|---|---|---|---|---|---|
| rvTWAS | AKAP13 | 15 | 85380571 | 85749358 | 8.87E-36 | - |
| rvTWAS | HBZ | 16 | 152687 | 154503 | 2.06E-21 | - |
| rvTWAS | LUC7L | 16 | 188969 | 229463 | 7.61E-12 | - |
| rvTWAS | PKMYT1 | 16 | 2968384 | 2980539 | 8.24E-96 | - |
| rvTWAS | PAQR4 | 16 | 2969245 | 2973489 | 1.48E-36 | - |
| rvTWAS | PDXDC1 | 16 | 14974591 | 15139339 | 1.58E-112 | - |
| rvTWAS | GDE1 | 16 | 19501890 | 19522145 | 3.12E-19 | - |
| rvTWAS | TBX6 | 16 | 30085793 | 30091887 | 4.03E-08 | - |
| rvTWAS | 01-Sep | 16 | 30378210 | 30395991 | 1.20E-55 | - |
| rvTWAS | RP11-212I21.4 | 16 | 55538200 | 55541837 | 1.10E-144 | - |
| rvTWAS | BBS2 | 16 | 56466836 | 56520283 | 2.09E-229 | - |
| rvTWAS | CLEC18A | 16 | 69950705 | 69964347 | 3.16E-08 | - |
| rvTWAS | CLEC18C | 16 | 70173449 | 70187256 | 1.01E-08 | - |
| rvTWAS | FUK | 16 | 70454421 | 70480274 | 1.05E-197 | - |
| rvTWAS | HP | 16 | 72054592 | 72061055 | 2.79E-07 | - |
| rvTWAS | HPR | 16 | 72063224 | 72077246 | 2.69E-07 | - |
| rvTWAS | APRT | 16 | 88809339 | 88811944 | 1.59E-20 | - |
| rvTWAS | ACADVL | 17 | 7217716 | 7225273 | 1.30E-10 | - |
| rvTWAS | EIF1 | 17 | 41688887 | 41692668 | 6.16E-16 | - |
| rvTWAS | NME1 | 17 | 51153536 | 51162428 | 7.50E-10 | - |
| rvTWAS | NME2 | 17 | 51165435 | 51171747 | 9.14E-08 | - |
| rvTWAS | NT5C | 17 | 75130266 | 75130867 | 7.65E-86 | - |
| rvTWAS | TNRC6C-AS1 | 17 | 78109051 | 78110457 | 2.91E-15 | - |
| rvTWAS | TPGS1 | 19 | 507834 | 519654 | 6.50E-37 | - |
| rvTWAS | HMG20B | 19 | 3572777 | 3579088 | 2.69E-120 | - |
| rvTWAS | CLPP | 19 | 6361452 | 6368908 | 1.29E-204 | - |
| rvTWAS | CD320 | 19 | 8302127 | 8308356 | 1.38E-183 | - |
| rvTWAS | CTD-2006C1.2 | 19 | 11987617 | 12046275 | 2.90E-15 | - |
| rvTWAS | SIN3B | 19 | 16829400 | 16880353 | 1.70E-94 | - |
| rvTWAS | RPL18A | 19 | 17859876 | 17864153 | 4.05E-06 | - |
| rvTWAS | BLOC1S3 | 19 | 45178745 | 45181801 | 1.23E-17 | - |
| rvTWAS | LILRA4 | 19 | 54333185 | 54339150 | 5.63E-63 | - |
| rvTWAS | LILRB4 | 19 | 54643889 | 54670359 | 5.27E-43 | - |
| rvTWAS | KIR2DL3 | 19 | 54738515 | 54753052 | 7.39E-12 | - |
| rvTWAS | KIR2DL1 | 19 | 54769811 | 54784322 | 1.33E-22 | - |
| rvTWAS | TGM2 | 20 | 38127781 | 38166578 | 3.17E-24 | + |
| rvTWAS | NCOA3 | 20 | 47501902 | 47656877 | 5.18E-11 | - |
| rvTWAS | UBE2V1 | 20 | 50081124 | 50115959 | 2.28E-40 | - |
| rvTWAS | FAM65C | 20 | 50586108 | 50691528 | 6.98E-11 | - |
| rvTWAS | BIRC7 | 20 | 63235883 | 63240507 | 1.66E-66 | - |

| | | | | | | |
|---|---|---|---|---|---|---|
| rvTWAS | IFNAR1 | 21 | 33324477 | 33359862 | 2.21E-45 | - |
| rvTWAS | TRPM2 | 21 | 44350163 | 44443081 | 9.54E-07 | + |
| rvTWAS | IGLV5-48 | 22 | 22352940 | 22353433 | 8.97E-09 | - |
| rvTWAS | CHCHD10 | 22 | 23765862 | 23768443 | 5.71E-21 | - |
| rvTWAS | NEFH | 22 | 29480243 | 29491390 | 1.67E-56 | - |
| rvTWAS | CYTH4 | 22 | 37282027 | 37315345 | 2.05E-09 | + |
| rvTWAS | EIF3L | 22 | 37849328 | 37876011 | 8.46E-21 | - |
| rvTWAS | TSPO | 22 | 43151514 | 43163242 | 3.44E-211 | + |
| rvTWAS | HDAC10 | 22 | 50245183 | 50249859 | 1.59E-06 | - |
| PrediXcan | EXT1 | 8 | 117794490 | 118111853 | 1.38E-21 | - |
| PrediXcan | APIP | 11 | 34853094 | 34916499 | 3.05E-18 | - |
| PrediXcan | OTX1 | 2 | 63050057 | 63057836 | 1.23E-16 | - |
| PrediXcan | RAB3GAP2 | 1 | 220148293 | 220272454 | 8.64E-16 | - |
| PrediXcan | IARS2 | 1 | 220094102 | 220148041 | 4.29E-14 | - |
| PrediXcan | SPINT1 | 15 | 40844018 | 40858207 | 7.38E-13 | - |
| PrediXcan | RNMT | 18 | 13726664 | 13764558 | 7.15E-12 | - |
| PrediXcan | CRYBA4 | 22 | 26621964 | 26630672 | 7.23E-12 | - |
| PrediXcan | PMM2 | 16 | 8788823 | 8848104 | 1.06E-11 | - |
| PrediXcan | ERICH6-AS1 | 3 | 150704043 | 150720146 | 2.78E-11 | - |
| PrediXcan | BRI3BP | 12 | 124993700 | 125031231 | 3.11E-11 | - |
| PrediXcan | DNAJB6 | 7 | 157335381 | 157417439 | 3.47E-11 | - |
| PrediXcan | SEMA3C | 7 | 80742538 | 80922359 | 3.94E-11 | - |
| PrediXcan | CHMP6 | 17 | 80991598 | 81009517 | 4.15E-11 | - |
| PrediXcan | ZNF350 | 19 | 51964343 | 51986856 | 4.19E-11 | - |
| PrediXcan | SPATA7 | 14 | 88384924 | 88470350 | 9.78E-11 | - |
| PrediXcan | RN7SL364P | 19 | 46688797 | 46689068 | 1.40E-10 | - |
| PrediXcan | SORD | 15 | 45023104 | 45077185 | 1.04E-09 | - |
| PrediXcan | TAAR1 | 6 | 132644984 | 132646003 | 1.05E-09 | - |
| PrediXcan | RP11-77H9.2 | 16 | 8849332 | 8860456 | 1.21E-09 | - |
| PrediXcan | FBXO2 | 1 | 11648367 | 11655785 | 1.79E-09 | - |
| PrediXcan | LARS | 5 | 146113038 | 146182660 | 2.39E-09 | - |
| PrediXcan | HIF1AN | 10 | 100529072 | 100559998 | 4.56E-09 | - |
| PrediXcan | CD36 | 7 | 80369575 | 80677008 | 4.76E-09 | + |
| PrediXcan | ANXA11 | 10 | 80150889 | 80205572 | 5.46E-09 | - |
| PrediXcan | RRBP1 | 20 | 17613678 | 17682295 | 5.83E-09 | - |
| PrediXcan | VENTX | 10 | 133237404 | 133241929 | 9.23E-09 | - |
| PrediXcan | DUSP13 | 10 | 75094432 | 75109221 | 1.44E-08 | - |
| PrediXcan | CARHSP1 | 16 | 8852942 | 8869012 | 1.59E-08 | - |
| PrediXcan | CYB561D2 | 3 | 50365334 | 50368137 | 1.84E-08 | - |
| PrediXcan | SCN2B | 11 | 118161951 | 118176673 | 2.10E-08 | - |
| PrediXcan | IL12RB1 | 19 | 18058995 | 18098944 | 2.28E-08 | - |
| PrediXcan | P3H2 | 3 | 189956728 | 190122437 | 2.41E-08 | - |

| | | | | | | |
|---|---|---|---|---|---|---|
| PrediXcan | CLDND1 | 3 | 98500840 | 98523066 | 9.27E-08 | - |
| PrediXcan | TMEM40 | 3 | 12733525 | 12769457 | 1.67E-07 | - |
| PrediXcan | RNF114 | 20 | 49936336 | 49953892 | 1.73E-07 | - |
| PrediXcan | MAST3 | 19 | 18097793 | 18151692 | 1.73E-07 | - |
| PrediXcan | ATG2B | 14 | 96279202 | 96363451 | 2.05E-07 | - |
| PrediXcan | PDK1 | 2 | 172555373 | 172608669 | 2.08E-07 | - |
| PrediXcan | SLC39A1 | 1 | 153959099 | 153967533 | 3.53E-07 | - |
| PrediXcan | ZFAND2A | 7 | 1152071 | 1160759 | 4.48E-07 | - |
| PrediXcan | PDLIM3 | 4 | 185500660 | 185535612 | 4.75E-07 | - |
| PrediXcan | NADK2 | 5 | 36192592 | 36242279 | 5.86E-07 | - |
| PrediXcan | SEPSECS-AS1 | 4 | 25160641 | 25201440 | 1.32E-06 | - |
| PrediXcan | LILRB2 | 19 | 54273821 | 54281184 | 1.37E-06 | - |
| PrediXcan | FANCM | 14 | 45135940 | 45200890 | 1.47E-06 | - |
| PrediXcan | INO80B | 2 | 74455023 | 74457960 | 1.62E-06 | - |
| PrediXcan | WDR41 | 5 | 77425970 | 77620611 | 1.69E-06 | - |
| PrediXcan | TMC8 | 17 | 78130770 | 78142968 | 2.72E-06 | - |
| PrediXcan | TUBGCP2 | 10 | 133278630 | 133311715 | 2.83E-06 | - |
| PrediXcan | LINC01146 | 14 | 88024550 | 88097619 | 2.95E-06 | - |
| PrediXcan | ACAD11 | 3 | 132558144 | 132644896 | 3.40E-06 | - |
| PrediXcan | A2M-AS1 | 12 | 9065177 | 9068060 | 3.58E-06 | - |
| kTWAS | EXO5 | 1 | 40508741 | 40516556 | 3.23E-26 | - |
| kTWAS | HLA-DQB2 | 6 | 32756098 | 32763534 | 1.86E-26 | - |
| kTWAS | INPP5E | 9 | 136428619 | 136439823 | 7.07E-54 | - |
| kTWAS | NDOR1 | 9 | 137205695 | 137217009 | 9.13E-47 | - |
| kTWAS | ZNF384 | 12 | 6666477 | 6689510 | 3.44E-84 | - |
| kTWAS | AKAP13 | 15 | 85380571 | 85749358 | 3.53E-35 | - |
| kTWAS | NME2 | 17 | 51165435 | 51171747 | 7.41E-10 | - |
| kTWAS | DNAH17 | 17 | 78423697 | 78577394 | 1.30E-10 | - |
| mkTWAS | TACSTD2 | 1 | 58575423 | 58577773 | 7.40E-07 | - |
| mkTWAS | RP11-63G10.4 | 1 | 58715609 | 58771295 | 5.77E-07 | - |
| mkTWAS | ADAMTSL4 | 1 | 150549369 | 150560894 | 5.42E-15 | - |
| mkTWAS | SEMA6C | 1 | 151131685 | 151146664 | 4.00E-72 | - |
| mkTWAS | GTF3C3 | 2 | 196763032 | 196799725 | 8.24E-10 | - |
| mkTWAS | GLB1L | 2 | 219236670 | 219245454 | 5.09E-07 | - |
| mkTWAS | RP11-689P11.2 | 4 | 8482270 | 8512610 | 2.53E-13 | - |
| mkTWAS | RP11-1277A3.1 | 5 | 177611253 | 177619754 | 5.75E-07 | - |
| mkTWAS | FAM153A | 5 | 177707981 | 177783398 | 1.97E-10 | - |
| mkTWAS | NRM | 6 | 30688047 | 30691420 | 2.25E-12 | - |

| | | | | | | |
|---|---|---|---|---|---|---|
| mkTWAS | XXbac-BPG252P9.9 | 6 | 30723105 | 30723339 | 1.84E-09 | - |
| mkTWAS | LY6G6C | 6 | 31718648 | 31721845 | 9.86E-08 | - |
| mkTWAS | NOL6 | 9 | 33461441 | 33473930 | 5.62E-08 | - |
| mkTWAS | SNAPC4 | 9 | 136375577 | 136400168 | 2.75E-15 | - |
| mkTWAS | C9orf163 | 9 | 136483495 | 136486067 | 6.25E-30 | - |
| mkTWAS | RP11-119F19.5 | 10 | 79681973 | 79682996 | 1.63E-07 | - |
| mkTWAS | MRPL23 | 11 | 1947278 | 1972793 | 1.51E-08 | - |
| mkTWAS | TRIM6 | 11 | 5596109 | 5612958 | 6.26E-18 | - |
| mkTWAS | HPX | 11 | 6431049 | 6442617 | 3.59E-14 | - |
| mkTWAS | RIN1 | 11 | 66332062 | 66336840 | 4.33E-37 | - |
| mkTWAS | B4GAT1 | 11 | 66345372 | 66347692 | 4.15E-73 | - |
| mkTWAS | ZNF384 | 12 | 6666477 | 6689510 | 1.96E-82 | - |
| mkTWAS | EID3 | 12 | 104303739 | 104305205 | 1.59E-19 | - |
| mkTWAS | RP11-131L12.3 | 12 | 118428281 | 118428870 | 1.17E-13 | - |
| mkTWAS | RP11-468E2.11 | 14 | 24201612 | 24202811 | 3.89E-10 | - |
| mkTWAS | MPP5 | 14 | 67241109 | 67335819 | 1.84632233037745e-317 | - |
| mkTWAS | ATP6V1D | 14 | 67294371 | 67359996 | 1.03E-288 | - |
| mkTWAS | IGHV3-33 | 14 | 106359793 | 106360324 | 6.64E-11 | - |
| mkTWAS | AKAP13 | 15 | 85380571 | 85749358 | 4.64E-29 | - |
| mkTWAS | RP11-303E16.2 | 16 | 81030770 | 81031485 | 1.62E-21 | - |
| mkTWAS | ATMIN | 16 | 81035847 | 81047358 | 4.17E-47 | - |
| mkTWAS | C16orf46 | 16 | 81053497 | 81077267 | 5.94E-38 | - |
| mkTWAS | RP1-59D14.5 | 17 | 2375061 | 2377746 | 7.98E-07 | - |
| mkTWAS | NME1 | 17 | 51153536 | 51162428 | 1.05E-09 | - |
| mkTWAS | DAPK3 | 19 | 3958453 | 3971123 | 5.57E-62 | - |
| mkTWAS | PIAS4 | 19 | 4007646 | 4039386 | 1.95E-68 | - |
| mkTWAS | MAP2K2 | 19 | 4090321 | 4124129 | 1.95E-68 | - |
| mkTWAS | UHRF1 | 19 | 4903080 | 4962154 | 6.11E-59 | - |
| mkTWAS | RINL | 19 | 38867834 | 38878279 | 5.77E-18 | - |
| mkTWAS | EID2 | 19 | 39538705 | 39540333 | 7.58E-16 | - |
| mkTWAS | ZNF574 | 19 | 42068477 | 42081549 | 3.43E-24 | - |
| mkTWAS | POU2F2 | 19 | 42086110 | 42196585 | 1.13E-30 | - |
| mkTWAS | CTC-378H22.2 | 19 | 42152569 | 42157523 | 9.78E-21 | - |
| mkTWAS | CEACAM1 | 19 | 42507304 | 42561234 | 4.83E-35 | - |
| mkTWAS | NKPD1 | 19 | 45149750 | 45158737 | 1.23E-17 | - |

| | | | | | | |
|---|---|---|---|---|---|---|
| mkTWAS | BLOC1S3 | 19 | 45178745 | 45181801 | 3.95E-18 | - |
| mkTWAS | OPA3 | 19 | 45527427 | 45602212 | 8.22E-07 | - |
| mkTWAS | PLA2G4C | 19 | 48047843 | 48110817 | 1.19E-13 | - |
| mkTWAS | GRWD1 | 19 | 48445773 | 48457022 | 5.80E-157 | - |
| mkTWAS | SPHK2 | 19 | 48624133 | 48630029 | 5.46E-168 | - |
| mkTWAS | BCAT2 | 19 | 48795062 | 48811029 | 5.06E-166 | - |
| mkTWAS | PPP1R15A | 19 | 48872392 | 48876057 | 1.12E-35 | - |
| mkTWAS | SNRNP70 | 19 | 49085419 | 49108605 | 1.79E-214 | - |
| mkTWAS | PIH1D1 | 19 | 49446298 | 49453497 | 9.46E-187 | - |
| mkTWAS | ALDH16A1 | 19 | 49453169 | 49471048 | 1.76E-08 | - |
| mkTWAS | NOSIP | 19 | 49555711 | 49590262 | 1.59E-60 | - |
| mkTWAS | PRRG2 | 19 | 49580646 | 49591015 | 3.91E-123 | - |
| mkTWAS | RRAS | 19 | 49635292 | 49640201 | 2.48E-219 | - |
| mkTWAS | FUZ | 19 | 49807113 | 49817376 | 2.29E-38 | - |
| mkTWAS | LILRB5 | 19 | 54249431 | 54257301 | 1.65E-16 | - |
| mkTWAS | LILRA4 | 19 | 54333185 | 54339150 | 6.27E-18 | - |
| mkTWAS | LILRB4 | 19 | 54643889 | 54670359 | 7.69E-23 | - |
| mkTWAS | PES1 | 22 | 30576625 | 30606837 | 3.00E-29 | - |
| mkTWAS | APOL1 | 22 | 36253010 | 36267530 | 9.90E-13 | - |

**Supplementary Table 3. Susceptibility genes identified by rvTWAS, PrediXcan, kTWAS and mkTWAS, at a Bonferroni-corrected P < 0.05 for Autism Spectrum Disorder (ASD)**

| Model | Gene | Chr | Start | End | P-value | Validated |
|---|---|---|---|---|---|---|
| rvTWAS | DISP3 | 1 | 11479166 | 11537584 | 2.07E-07 | - |
| rvTWAS | NECAP2 | 1 | 16440672 | 16460078 | 1.50E-07 | - |
| rvTWAS | LINC01772 | 1 | 16460948 | 16463287 | 2.48E-07 | - |
| rvTWAS | NBPF1 | 1 | 16562319 | 16613562 | 1.02E-07 | - |
| rvTWAS | CR1L | 1 | 207645174 | 207738416 | 2.02E-06 | - |
| rvTWAS | TTN-AS1 | 2 | 178521183 | 178779963 | 2.28E-06 | - |
| rvTWAS | INPP5D | 2 | 233059967 | 233207903 | 3.36E-06 | - |
| rvTWAS | IL17RC | 3 | 9917074 | 9933621 | 1.90E-06 | - |
| rvTWAS | TRANK1 | 3 | 36826820 | 36945057 | 3.23E-06 | - |
| rvTWAS | ACSL6 | 5 | 131949973 | 132012243 | 3.34E-06 | - |
| rvTWAS | ANKIB1 | 7 | 92246234 | 92401384 | 2.28E-07 | - |
| rvTWAS | DNAJB5 | 9 | 34989641 | 34998900 | 3.30E-06 | - |
| rvTWAS | VCP | 9 | 35056064 | 35073249 | 4.53E-06 | + |
| rvTWAS | XXyac-YM21GA2.7 | 9 | 87857015 | 87859399 | 5.89E-06 | - |
| rvTWAS | DPYSL4 | 10 | 132186900 | 132205776 | 6.52E-06 | - |
| rvTWAS | SLC15A3 | 11 | 60937160 | 60952530 | 3.15E-08 | - |
| rvTWAS | RILPL1 | 12 | 123470054 | 123533718 | 3.97E-06 | - |
| rvTWAS | EIF2B1 | 12 | 123620942 | 123633738 | 1.44E-06 | - |
| rvTWAS | ATP6V0A2 | 12 | 123712318 | 123761755 | 4.78E-06 | - |
| rvTWAS | IGHV4-4 | 14 | 106011922 | 106012420 | 9.08E-07 | - |
| rvTWAS | AP3B2 | 15 | 82659281 | 82709914 | 1.98E-06 | + |
| rvTWAS | CMTR2 | 16 | 71281389 | 71289715 | 1.56E-06 | - |
| rvTWAS | NUP88 | 17 | 5360963 | 5419640 | 5.29E-06 | - |
| rvTWAS | CHD3 | 17 | 7884806 | 7912760 | 1.36E-07 | - |
| rvTWAS | MPP2 | 17 | 43875357 | 43909711 | 4.73E-06 | - |
| rvTWAS | ASB16 | 17 | 44170447 | 44179083 | 3.02E-07 | - |
| rvTWAS | ASB16-AS1 | 17 | 44175973 | 44186717 | 3.02E-07 | - |
| rvTWAS | EIF4A3 | 17 | 80135214 | 80147183 | 9.21E-09 | - |
| rvTWAS | CCDC57 | 17 | 82101460 | 82212830 | 3.34E-06 | - |
| rvTWAS | PPP4R1 | 18 | 9546791 | 9615240 | 5.07E-07 | - |
| rvTWAS | SLC39A3 | 19 | 2732204 | 2740152 | 2.80E-06 | - |
| rvTWAS | PIP5K1C | 19 | 3630183 | 3700479 | 1.89E-06 | - |
| rvTWAS | FEM1A | 19 | 4791681 | 4801273 | 8.07E-08 | - |
| rvTWAS | ZNF90 | 19 | 20077994 | 20127076 | 3.24E-06 | - |
| rvTWAS | MYPOP | 19 | 45890020 | 45902604 | 6.29E-06 | - |
| rvTWAS | C19orf48 | 19 | 50797704 | 50804929 | 2.41E-06 | - |
| rvTWAS | LLNLR-470E3.1 | 19 | 51639478 | 51639931 | 4.75E-07 | - |
| rvTWAS | SIGLEC14 | 19 | 51642553 | 51646801 | 2.95E-07 | - |
| rvTWAS | LILRA4 | 19 | 54333185 | 54339150 | 4.35E-06 | - |

| | | | | | | |
|---|---|---|---|---|---|---|
| rvTWAS | PIK3IP1 | 22 | 31281593 | 31292498 | 9.94E-07 | - |
| rvTWAS | CHKB | 22 | 50578949 | 50601455 | 2.61E-07 | - |
| PrediXcan | IQGAP3 | 1 | 156525405 | 156572604 | 5.89E-09 | + |
| PrediXcan | MPC2 | 1 | 167916729 | 167937040 | 9.80E-09 | - |
| PrediXcan | RP11-85K15.3 | 14 | 35086035 | 35088059 | 9.92E-09 | - |
| PrediXcan | RP11-117D22.2 | 1 | 53366668 | 53368245 | 1.28E-08 | - |
| PrediXcan | CTD-2574D22.4 | 16 | 29919865 | 29921905 | 1.85E-08 | - |
| PrediXcan | RP11-36D19.9 | 10 | 80249710 | 80254121 | 1.92E-08 | - |
| PrediXcan | ADCK1 | 14 | 77800083 | 77933954 | 2.09E-08 | - |
| PrediXcan | C1QBP | 17 | 5432877 | 5448808 | 2.18E-08 | - |
| PrediXcan | TCIRG1 | 11 | 68039016 | 68050664 | 3.78E-08 | - |
| PrediXcan | DOT1L | 19 | 2164149 | 2232578 | 4.36E-08 | - |
| PrediXcan | PPP2R3C | 14 | 35085467 | 35122517 | 6.27E-08 | - |
| PrediXcan | RP11-526I2.5 | 15 | 100547765 | 100550153 | 6.35E-08 | - |
| PrediXcan | NDUFAF7 | 2 | 37231714 | 37253403 | 7.37E-08 | - |
| PrediXcan | ERGIC1 | 5 | 172834275 | 172952685 | 8.39E-08 | - |
| PrediXcan | SERHL2 | 22 | 42553617 | 42572529 | 1.03E-07 | - |
| PrediXcan | RNASEH1-AS1 | 2 | 3558617 | 3561745 | 1.37E-07 | - |
| PrediXcan | SCLT1 | 4 | 128864921 | 129093316 | 2.42E-07 | - |
| PrediXcan | RUBCNL | 13 | 46342000 | 46438190 | 2.43E-07 | - |
| PrediXcan | SMPD4 | 2 | 130151392 | 130182626 | 2.66E-07 | - |
| PrediXcan | DNASE2 | 19 | 12875211 | 12881468 | 3.10E-07 | - |
| PrediXcan | MAEA | 4 | 1289851 | 1340137 | 3.63E-07 | - |
| PrediXcan | SPATA2L | 16 | 89696365 | 89701705 | 3.64E-07 | - |
| PrediXcan | RP11-298J23.10 | 6 | 41783606 | 41784118 | 4.16E-07 | - |
| PrediXcan | RP4-612B15.3 | 1 | 86703502 | 86704462 | 5.38E-07 | - |
| PrediXcan | CRABP2 | 1 | 156699606 | 156705816 | 5.98E-07 | - |
| PrediXcan | LMNA | 1 | 156082573 | 156140089 | 6.78E-07 | + |
| PrediXcan | TFEB | 6 | 41683978 | 41736259 | 8.82E-07 | - |
| PrediXcan | PACSIN2 | 22 | 42858387 | 43015145 | 1.01E-06 | - |
| PrediXcan | SLC8B1 | 12 | 113298759 | 113359493 | 1.32E-06 | - |
| PrediXcan | IGSF8 | 1 | 160091340 | 160098943 | 1.37E-06 | - |
| PrediXcan | ATG4B | 2 | 241637450 | 241673857 | 1.43E-06 | - |
| PrediXcan | LINC00672 | 17 | 38925168 | 38929384 | 1.58E-06 | - |
| PrediXcan | NEO1 | 15 | 73051710 | 73305206 | 1.63E-06 | + |
| PrediXcan | RBM23 | 14 | 22893206 | 22917802 | 1.84E-06 | - |
| PrediXcan | RP11-230F18.5 | 13 | 113527260 | 113530621 | 1.92E-06 | - |
| PrediXcan | RP11-38G5.2 | 15 | 79832466 | 79833554 | 1.92E-06 | - |
| PrediXcan | FAM114A2 | 5 | 153990128 | 154038905 | 2.13E-06 | - |
| PrediXcan | RP11-288C18.1 | 2 | 36839922 | 36842539 | 2.50E-06 | - |
| PrediXcan | GPNMB | 7 | 23235967 | 23275108 | 3.20E-06 | - |
| PrediXcan | CTD-2054N24.2 | 15 | 99807023 | 99877148 | 3.72E-06 | - |
| PrediXcan | STX10 | 19 | 13144058 | 13150383 | 4.13E-06 | - |

| | | | | | | |
|---|---|---|---|---|---|---|
| kTWAS | ST6GALNAC3 | 1 | 76074719 | 76634601 | 1.36E-06 | - |
| kTWAS | CR1L | 1 | 207645174 | 207738416 | 1.84E-06 | - |
| kTWAS | DNAJB5 | 9 | 34989641 | 34998900 | 2.44E-06 | - |
| kTWAS | SLC15A3 | 11 | 60937160 | 60952530 | 3.15E-08 | - |
| kTWAS | FEN1 | 11 | 61792803 | 61797244 | 2.01E-08 | - |
| kTWAS | RILPL1 | 12 | 123470054 | 123533718 | 4.14E-06 | - |
| kTWAS | ATP6V0A2 | 12 | 123712318 | 123761755 | 4.03E-06 | - |
| kTWAS | EP400NL | 12 | 132084283 | 132131639 | 1.61E-08 | - |
| kTWAS | CAPN3 | 15 | 42359500 | 42412317 | 1.57E-06 | - |
| kTWAS | AMFR | 16 | 56361452 | 56425538 | 2.54E-06 | - |
| kTWAS | ADAM11 | 17 | 44759031 | 44781846 | 3.15E-09 | - |
| kTWAS | ARL17A | 17 | 46516702 | 46579682 | 1.97E-06 | - |
| kTWAS | CCDC57 | 17 | 82101460 | 82212830 | 6.61E-06 | - |
| kTWAS | ANKRD24 | 19 | 4183354 | 4224814 | 2.93E-06 | - |
| kTWAS | ZNF506 | 19 | 19785839 | 19821751 | 1.29E-06 | - |
| kTWAS | C19orf48 | 19 | 50797704 | 50804929 | 2.43E-06 | - |
| kTWAS | EYA2 | 20 | 46894624 | 47188844 | 2.99E-06 | - |
| kTWAS | NDUFV3 | 21 | 42879644 | 42913304 | 6.05E-06 | - |
| kTWAS | PIK3IP1 | 22 | 31281593 | 31292498 | 2.65E-06 | - |
| mkTWAS | PDZK1IP1 | 1 | 47183593 | 47191044 | 2.87E-08 | - |
| mkTWAS | TAL1 | 1 | 47216290 | 47232220 | 1.21E-10 | + |
| mkTWAS | TTN-AS1 | 2 | 178521183 | 178779963 | 3.80E-07 | - |
| mkTWAS | IL17RC | 3 | 9917074 | 9933621 | 2.48E-06 | - |
| mkTWAS | RASGEF1B | 4 | 81426393 | 82044244 | 7.49E-07 | - |
| mkTWAS | ENOPH1 | 4 | 82430562 | 82461091 | 1.04E-07 | - |
| mkTWAS | SPEF2 | 5 | 35617844 | 35814611 | 4.44E-07 | - |
| mkTWAS | VCP | 9 | 35056064 | 35073249 | 2.23E-06 | + |
| mkTWAS | RP11-297B17.3 | 9 | 37002777 | 37008040 | 3.61E-07 | - |
| mkTWAS | EIF2B1 | 12 | 123620942 | 123633738 | 1.88E-06 | - |
| mkTWAS | C14orf79 | 14 | 104985775 | 104999527 | 2.15E-07 | - |
| mkTWAS | LA16c-390H2.4 | 16 | 3542776 | 3545785 | 1.48E-06 | - |
| mkTWAS | CMTR2 | 16 | 71281389 | 71289715 | 1.37E-06 | - |
| mkTWAS | SENP3 | 17 | 7561875 | 7571969 | 3.13E-09 | - |
| mkTWAS | DRC3 | 17 | 17972813 | 18016889 | 1.92E-06 | - |
| mkTWAS | G6PC3 | 17 | 44070735 | 44076344 | 6.50E-09 | - |
| mkTWAS | C17orf53 | 17 | 44141906 | 44162476 | 3.27E-09 | - |
| mkTWAS | UBTF | 17 | 44205033 | 44221626 | 1.15E-08 | - |
| mkTWAS | ACBD4 | 17 | 45132600 | 45144181 | 4.16E-07 | - |
| mkTWAS | RP11-855A2.2 | 17 | 67973934 | 67976072 | 4.94E-07 | - |
| mkTWAS | RP11-143J12.3 | 18 | 9112404 | 9115877 | 4.67E-07 | - |
| mkTWAS | DIRAS1 | 19 | 2714567 | 2721418 | 1.61E-06 | - |
| mkTWAS | ZNF555 | 19 | 2841435 | 2860484 | 5.49E-07 | - |
| mkTWAS | ZNF506 | 19 | 19785839 | 19821751 | 1.29E-06 | - |

| | | | | | | |
|---|---|---|---|---|---|---|
| mkTWAS | ZNF90 | 19 | 20077994 | 20127076 | 2.29E-07 | - |
| mkTWAS | CTC-260E6.6 | 19 | 20150361 | 20321305 | 5.02E-07 | - |
| mkTWAS | CTC-260E6.4 | 19 | 20220597 | 20221868 | 7.14E-07 | - |
| mkTWAS | SPATA25 | 20 | 45886489 | 45887635 | 8.79E-08 | - |
| mkTWAS | FAM209B | 20 | 56533246 | 56536520 | 2.24E-06 | - |

#### Supplementary Figures

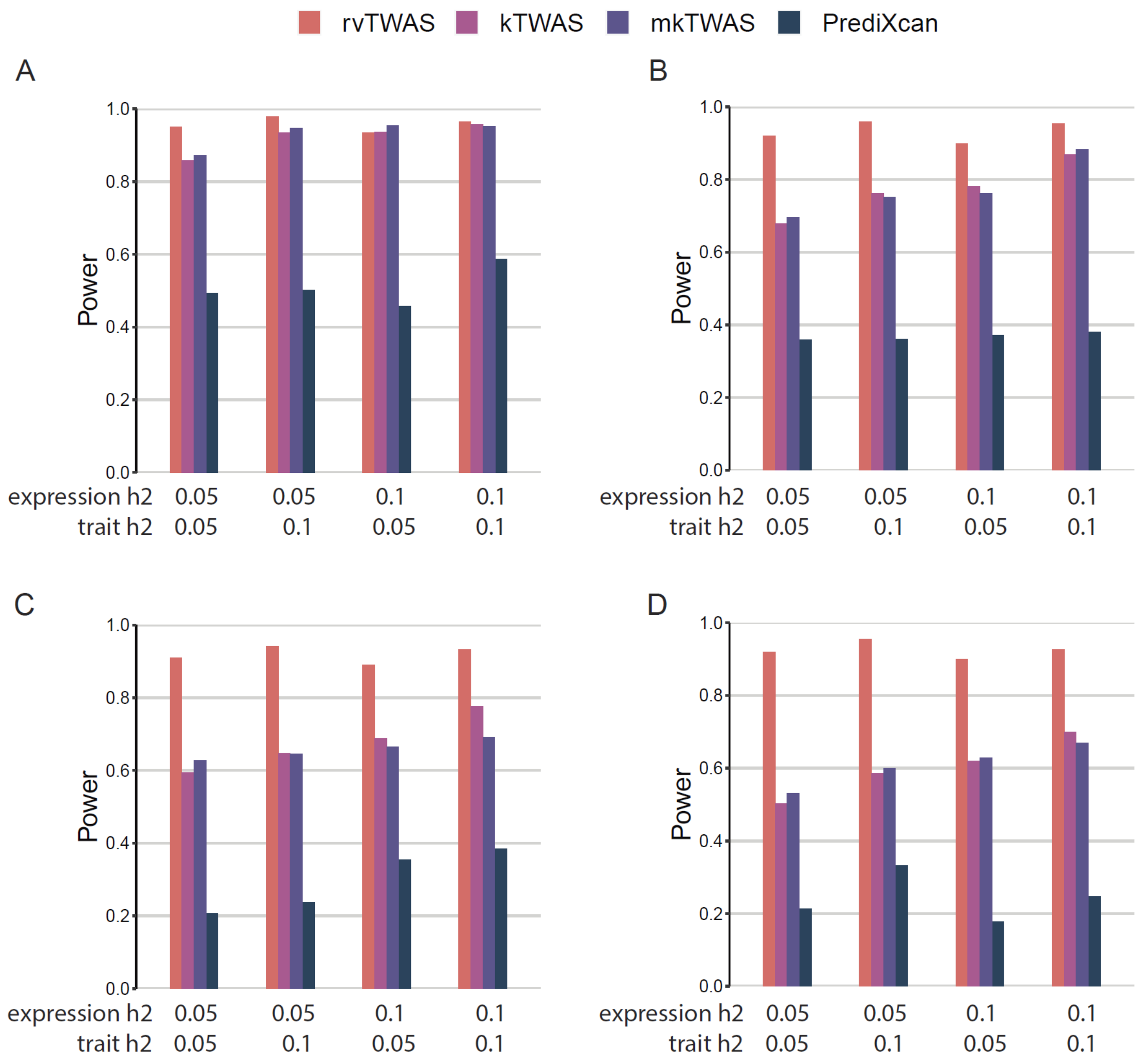

**Supplementary Figure 1. Power comparison under pleiotropy scenarios.** Power is indicated on the y-axis. All panels are results under an additive genetic architecture (Sum), with differing expression heritability and trait heritability denoted below each panel. The total number of contributing rare genetic variants (with MAF < 0.005) M are fixed in each panel. **A**) M=0; **B**) M=50; **C**) M=100; and **D**) M=200.

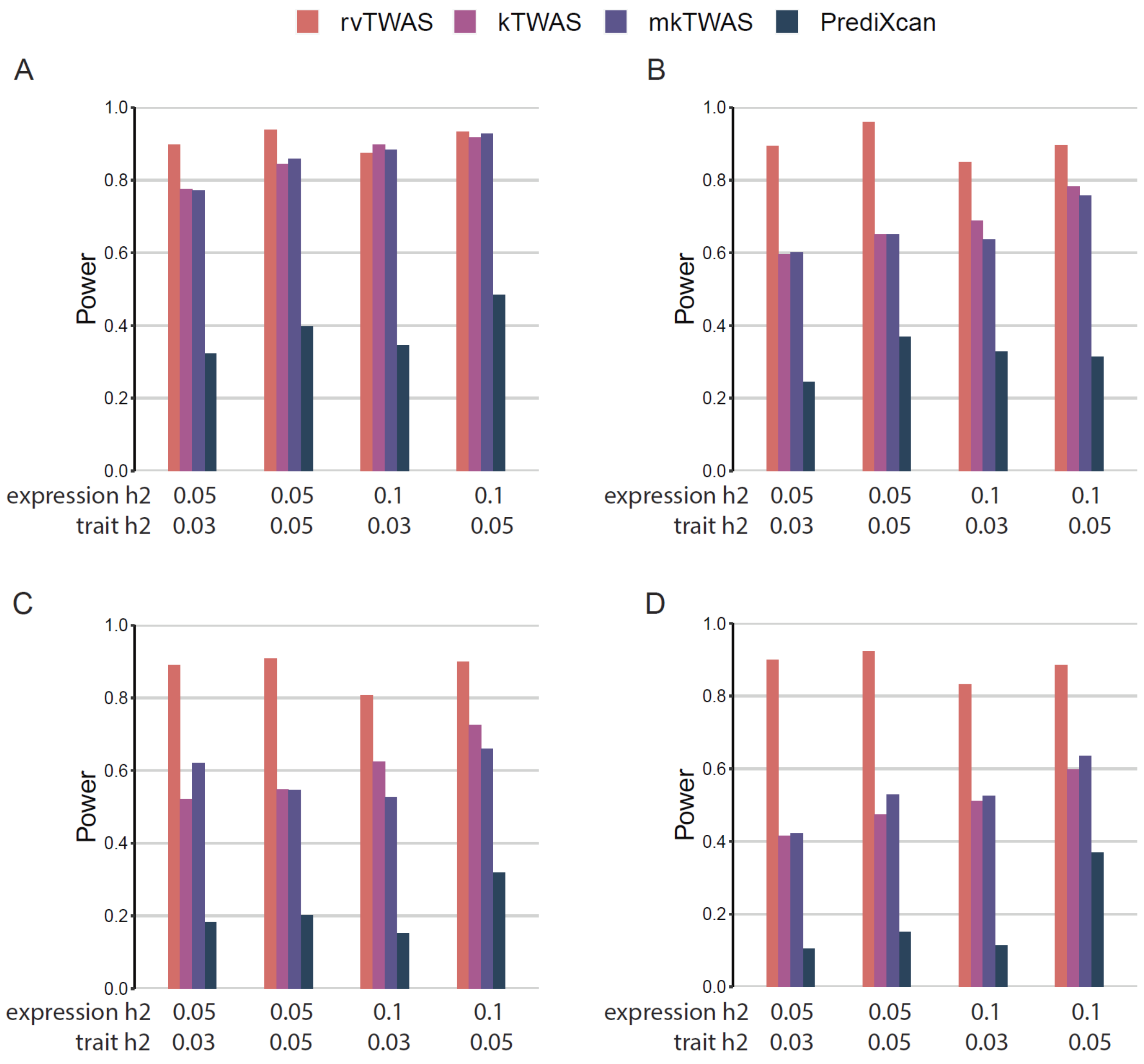

**Supplementary Figure 2. Power comparison under causality scenarios.** Power is indicated on the y-axis. All panels are results under an additive genetic architecture (Sum), with differing expression heritability and trait heritability denoted below each panel. The total number of contributing rare genetic variants (with MAF < 0.005) M are fixed in each panel. **A**) M=0; **B**) M=50; **C**) M=100, and **D**) M=200.

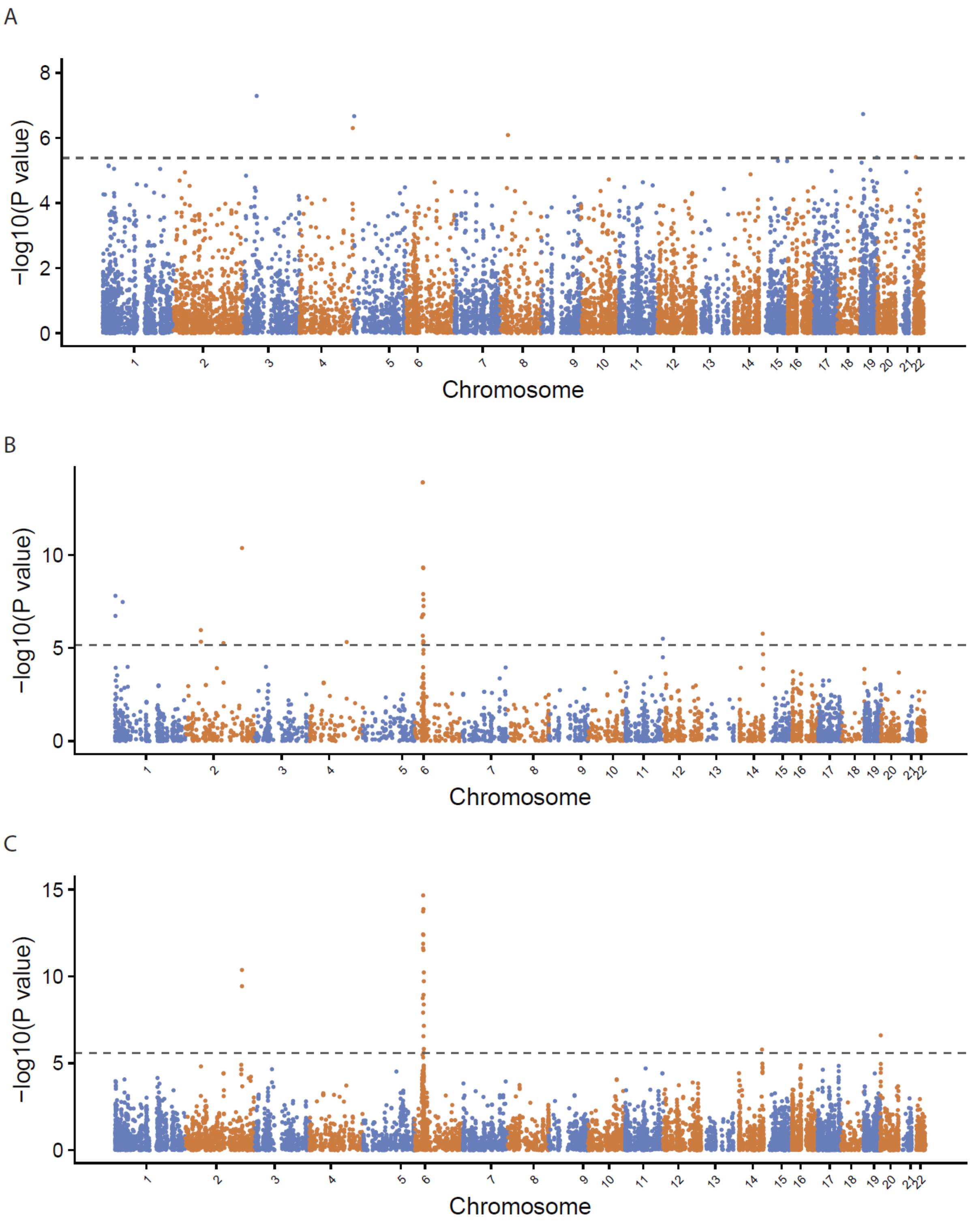

**Supplementary Figure 3. Manhattan plots showing associations identified by PrediXcan, kTWAS and mkTWAS for schizophrenia (SCZ).** Black dashed lines indicate the cut-off specified by Bonferroni correction at the level of P-value < 0.05. A) PrediXcan. B) kTWAS. C) mkTWAS.

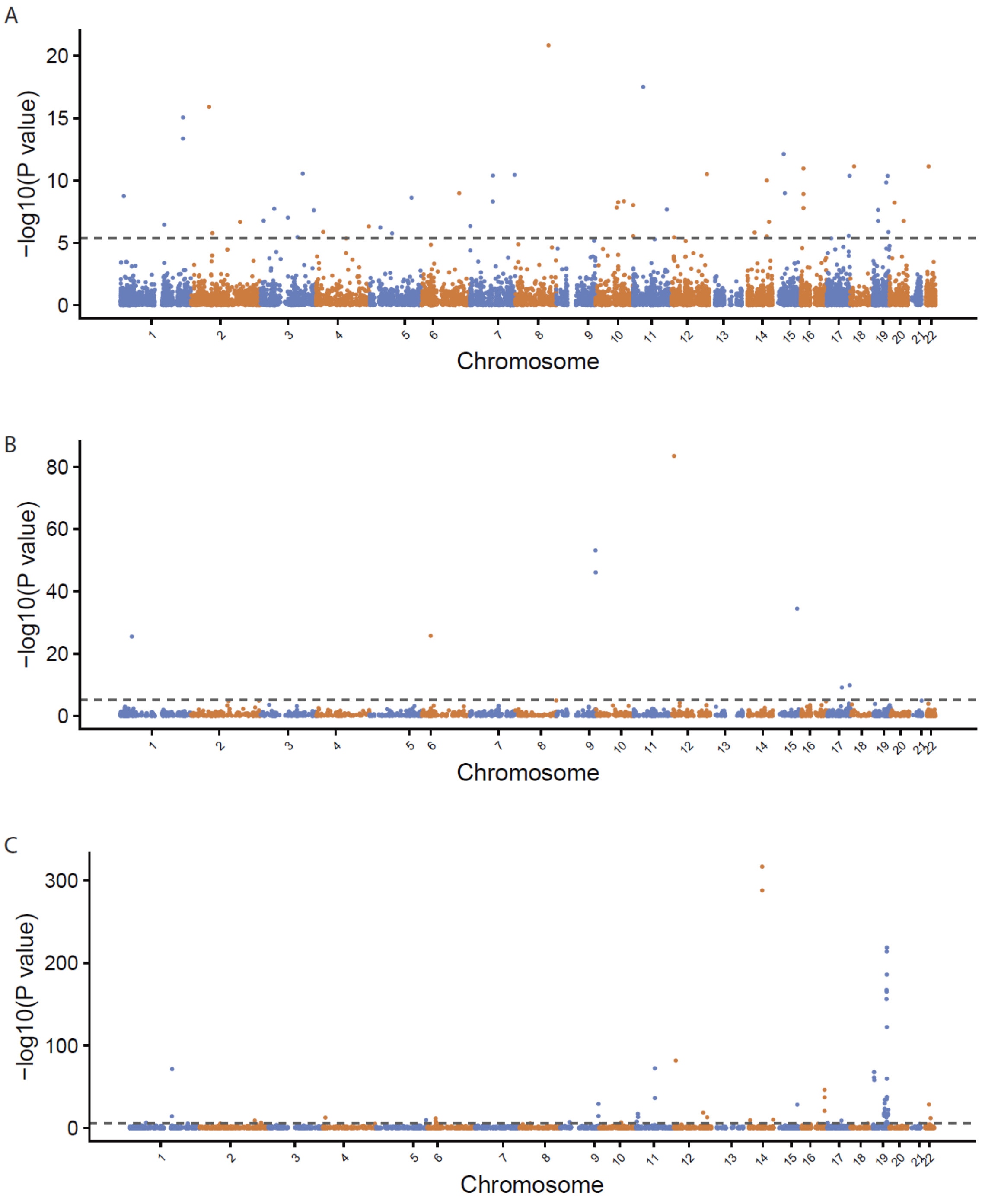

**Supplementary Figure 4. Manhattan plots showing associations identified by PrediXcan, kTWAS and mkTWAS for bipolar disorder (BPD).** Black dashed lines indicate the cut-off specified by Bonferroni correction at the level of P-value < 0.05. A) PrediXcan. B) kTWAS. C) mkTWAS.

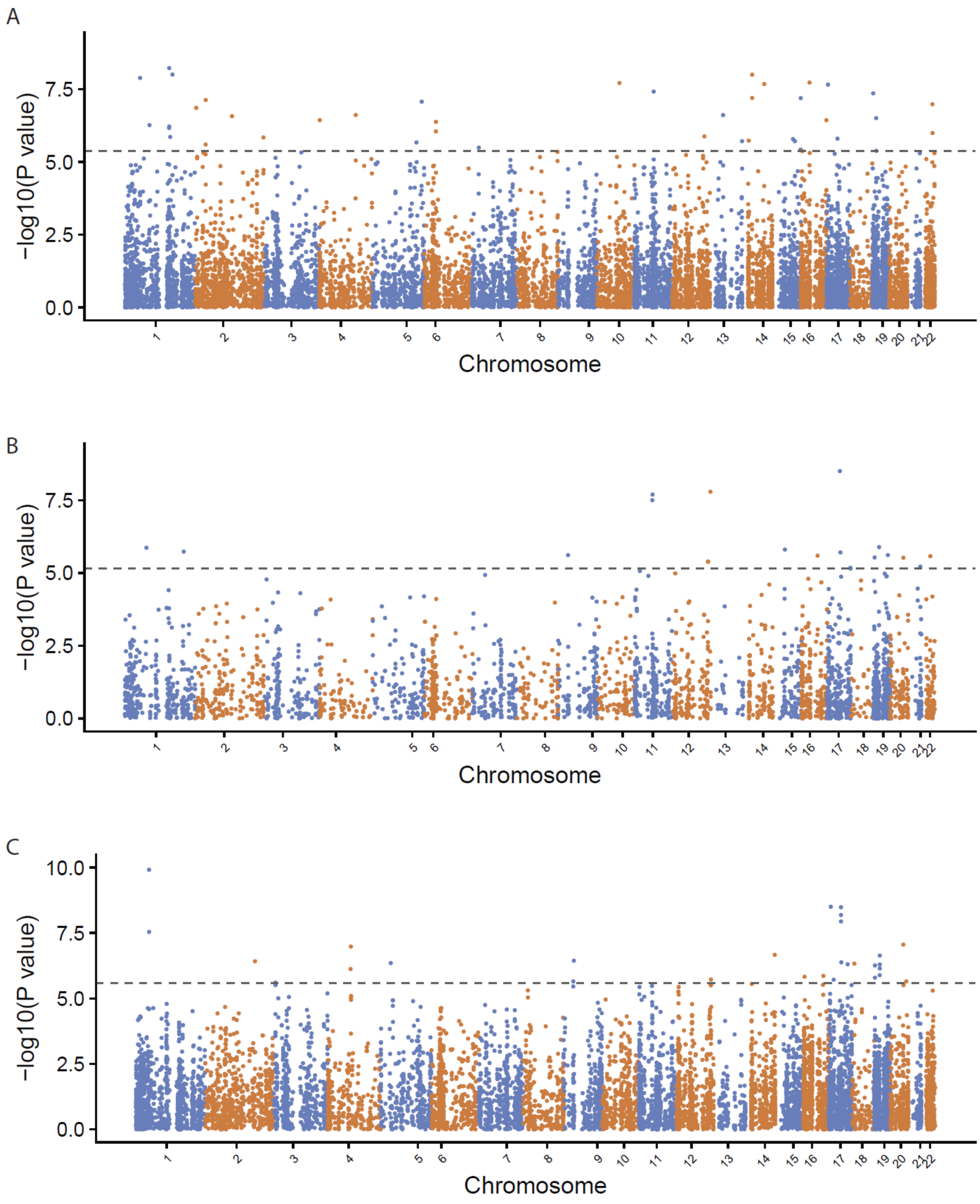

**Supplementary Figure 5. Manhattan plots showing associations identified by PrediXcan, kTWAS and mkTWAS for autism spectrum disorder (ASD).** Black dashed lines indicate the cut-off specified by Bonferroni correction at the level of P-value < 0.05. A) PrediXcan. B) kTWAS. C) mkTWAS.

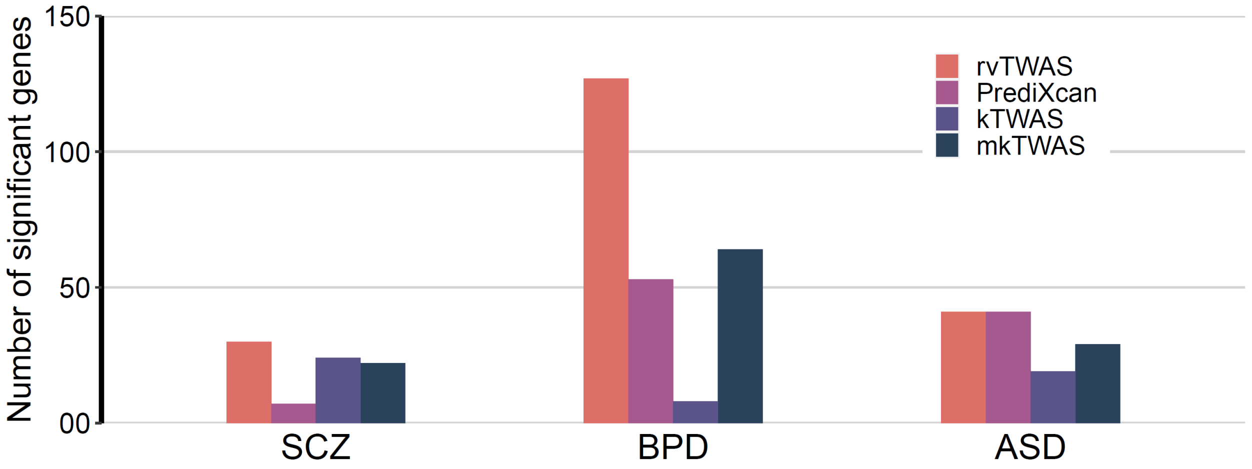

**Supplementary Figure 6. The comparisons of rvTWAS and other TWAS methods for SCZ, BPD and ASD.** The number of significant genes discovered by rvTWAS, PrediXcan, kTWAS, and mkTWAS under Bonferroni corrected *P*<0.05. The left, middle, and right columns are for SCZ, BPD and ASD, respectively.
